# Malonyl-CoA is an ancient physiological ATP-competitive mTORC1 inhibitor

**DOI:** 10.1101/2022.11.06.515351

**Authors:** Raffaele Nicastro, Laura Brohée, Josephine Alba, Julian Nüchel, Gianluca Figlia, Stefanie Kipschull, Peter Gollwitzer, Jesus Romero-Pozuelo, Stephanie A. Fernandes, Andreas Lamprakis, Stefano Vanni, Aurelio A. Teleman, Claudio De Virgilio, Constantinos Demetriades

**Author notes:** Correspondence and requests for materials should be addressed to Constantinos Demetriades, Claudio De Virgilio, Aurelio Teleman, or Stefano Vanni. These authors contributed equally to this work (Raffaele Nicastro, Laura Brohée). These authors jointly supervised this work (Stefano Vanni, Aurelio A. Teleman, Claudio De Virgilio, Constantinos Demetriades). Center for Biological Research (CIB-CSIC), 28040 Madrid, Spain.

## Abstract

Cell growth is regulated primarily by the mammalian/mechanistic Target of Rapamycin Complex 1 (mTORC1) that functions both as a nutrient sensor and a master controller of virtually all biosynthetic pathways ^1^. This ensures that cells are metabolically active only when conditions are optimal for growth. Notably, although mTORC1 is known to regulate fatty acid (FA) biosynthesis, how and whether the cellular lipid biosynthetic capacity signals back to fine-tune mTORC1 activity remains poorly understood. Here, we show that mTORC1 senses the capacity of a cell to synthesize FAs by detecting the levels of malonyl-CoA, an intermediate of this biosynthetic pathway. We find that, in both yeast and mammalian cells, this regulation is very direct, with malonyl-CoA binding to the mTOR catalytic pocket and acting as a specific ATP-competitive inhibitor. When ACC1 (acetyl-CoA carboxylase 1) is hyperactive or FASN (fatty acid synthase) is downregulated/inhibited, elevated malonyl-CoA levels are channelled to proximal mTOR molecules that form direct protein-protein interactions with ACC1 and FASN. Our findings represent a conserved, unique, homeostatic mechanism whereby impaired FA biogenesis leads to reduced mTORC1 activity to coordinatively link this metabolic pathway to the overall cellular biosynthetic output. Moreover, they reveal the first-described example of a physiological metabolite that directly inhibits the activity of a signalling kinase by competing with ATP for binding.

---

Cell growth is a highly energy-consuming and, hence, tightly-regulated process. Cells accumulate mass by taking up essential nutrients from their environment and using them to build macromolecules, such as proteins, lipids, and sugars, through intricate metabolic pathways ^2^. mTORC1 is a central integration point in cellular signalling, linking metabolic cues to cell growth and homeostasis. Work over the last 15 years has identified complex signalling cascades through which growth factors (GFs), nutrients (like amino acids; AAs), and energy availability regulate mTORC1 (reviewed in ^1-4^). Accordingly, cholesterol levels were previously shown to influence mTORC1 activity at the lysosomal surface via mechanisms that involve the Niemann-Pick C1 (NPC1), SLC38A9 and Rag GTPase proteins ^5-7^, with the latter also playing a central role in AA and glucose sensing by recruiting mTOR at this subcellular location ^8-10^. In turn, mTORC1 upregulates multiple metabolic processes, including protein and cholesterol biosynthesis ^3,4^. Similarly, active mTORC1 promotes FA biosynthesis by driving the expression of key enzymes in this process—like FASN—in response to GF signalling ^11^. Interestingly, however, whether and how FA synthesis controls mTORC1 activity remains enigmatic.

Fatty acids serve both structural and regulatory roles in cells (e.g., by participating in membrane formation and by post-translationally modifying proteins, respectively), while also functioning as energy storage molecules ^12,13^. In *de novo* FA biosynthesis, acetyl-CoA, generated from glucose catabolism, is converted to malonyl-CoA by ACC1-mediated, ATP-dependent carboxylation, which is then used by FASN to produce palmitate, the precursor of longer fatty acids ^14,15^. This process is well conserved from human to yeast cells, where the homologous acetyl-CoA carboxylase, Acc1, and the Fas1- and Fas2-containing fatty acid synthase (FAS) complex catalyse the conversion of acetyl-CoA to FAs ^16^ (Fig. 1a). Because various metabolites were previously shown to impact mTORC1 via regulating the activity of upstream signalling components (e.g., by AMP and inositol allosterically influencing AMPK activity ^17,18^; or α-ketoglutarate and glutaminolysis somehow affecting the lysosomal Rag GTPases ^19^), we hypothesized that a metabolic intermediate of FA biosynthesis may be signalling directly or indirectly to mTORC1, thus forming a regulatory feedback loop.

**Figure 1.**
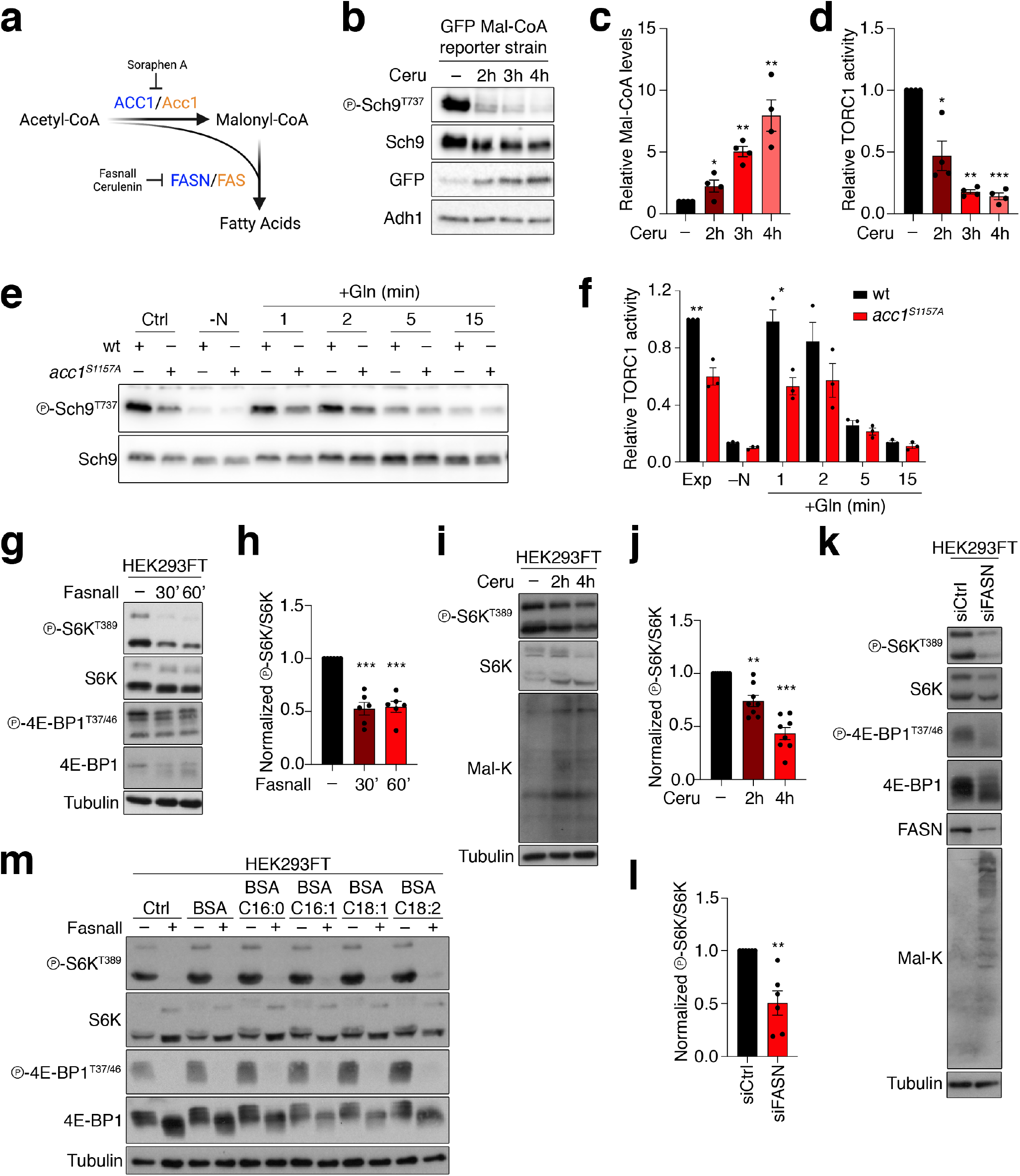
Increasing malonyl-CoA levels by genetic and pharmacological perturbation of Fas1/Acc1 activity reduces mTORC1 activity in yeast and mammalian cells. **(a)** Schematic diagram of *de novo* fatty acid biosynthesis. Yeast and mammalian proteins shown in orange or blue, respectively. **(b-d)** Cerulenin boosts malonyl-CoA levels and inhibits TORC1 activity. Immunoblots with lysates from yeast cells expressing the fapR/fapOp-yeGFP malonyl-CoA reporter system, treated with 20 μM cerulenin (Ceru) for the indicated times. GFP expression shows Mal-CoA levels. TORC1 activity assessed by Sch9 phosphorylation (b). Quantification of relative Mal-CoA levels (GFP/Adh1 ratio) in (c). Quantification of relative TORC1 activity (p-Sch9^T737^/Sch9 ratio) in (d), n = 4. **(e-f)** Immunoblots with lysates from wild-type (wt) or Acc1^S1157A^-expressing yeast cells, exponentially growing (Ctrl), starved for nitrogen (-N), or starved and restimulated with Gln for the indicated times. TORC1 activity assessed by Sch9 phosphorylation (e). Quantification of relative TORC1 activity (p-Sch9^T737^/Sch9 ratio) in (f), n = 3. **(g-h)** Pharmacological inhibition of FASN downregulates mTORC1. Immunoblots with lysates from HEK293FT cells treated with 25 μM Fasnall for the indicated times. mTORC1 activity assayed by phosphorylation of S6K and 4E-BP1 (g). Quantification of p-S6K^T389^/S6K ratio in (h), n = 6. **(i-j)** Immunoblots with lysates from HEK293FT cells treated with 50 μM Cerulenin (Ceru) for the indicated times. mTORC1 activity assayed by phosphorylation of S6K. Malonyl-lysine (Mal-K) blots indicate total protein malonylation, as a read-out for Mal-CoA levels (i). Quantification of p-S6K^T389^/S6K ratio in (j), n = 8. **(k-l)** Immunoblots with lysates from control (siCtrl) or FASN knockdown (siFASN) HEK293FT cells. mTORC1 activity assayed by phosphorylation of S6K and 4E-BP1. Mal-K blots show total protein malonylation (k). Quantification of p-S6K^T389^/S6K ratio in (l), n = 6. **(m)** FASN inhibition downregulates mTORC1 activity independently from lipid availability. Immunoblots with lysates from control (–) or Fasnall-treated (25 μM, 30 min) HEK293FT cells, supplemented with BSA-conjugated FAs as indicated, or BSA as control. mTORC1 activity assayed by phosphorylation of S6K and 4E-BP1. Data in graphs shown as mean ± SEM. * = p < 0.05, ** = p < 0.005, *** = p < 0.0005.

## Functional pharmacogenetic interactions between TOR pathway components and the core FA biosynthesis machinery reveal a role for malonyl-CoA in TOR signalling

Our previous interactome studies of the Rag GTPases in yeast indicated that the RagA/B homologue, Gtr1, may directly or indirectly interact with Acc1 and the Fas1 subunit of the FAS complex (see Supplemental Table S1 in ^20^). Because both of these enzymes are part of essential protein complexes that work together to synthesize FAs, this suggested a possible functional interplay between TORC1 signalling and the core FA biosynthetic machinery. To study this, we first probed the pharmacogenetic interaction of TORC1 pathway mutants with the FAS inhibitor cerulenin ^21,22^. Interestingly, loss of Gtr1/2, expression of the nucleotide-free Gtr1^S20L^ allele, loss of other TORC1 pathway components (i.e., Ego1-3, Tor1, Tco89), or co-treatment with rapamycin, all of which reduce TORC1 activity ^23^, rendered cells highly sensitive to cerulenin (Extended Data Fig. 1a,b). Conversely, expression of the GTP-locked Gtr1^Q65L^ allele, which activates TORC1, caused cerulenin resistance (Extended Data Fig. 1a). These observations match our targeted re-analyses of recently published SATAY (saturated transposon analyses in yeast) data ^24,25^, showing that transposon events in TORC1 pathway genes were under-represented when cells were grown on cerulenin or rapamycin (Extended Data Fig. 1c,e). Intriguingly, however, in these datasets, TORC1 pathway genes were over-represented when cells were grown in the presence of soraphen A, which inhibits the antecedent enzyme in FA biosynthesis, Acc1 ^26^ (Extended Data Fig. 1d,e). One interpretation of these findings is that malonyl-CoA, the intermediate metabolite between Acc1 and FAS, may be functionally linked to TORC1 activity. In support of this hypothesis, cerulenin-mediated FAS inhibition, which increases intracellular malonyl-CoA levels more than 8-fold (as assayed with a specific reporter system ^27^) (Fig. 1b,c), strongly reduced TORC1 activity (assayed by the phosphorylation of its direct target Sch9) *in vivo* (Fig. 1b,d). Next, we hypothesised that FAS inhibition could be downregulating TORC1 either by increasing the levels of its substrate (i.e., malonyl-CoA) or by reducing the levels of its product (i.e., palmitate). To distinguish between these two possibilities, we used a hyperactive Acc1^S1157A^ allele, which causes malonyl-CoA levels to increase (Extended Data Fig. 2c,d) ^27-29^. This allele aggravated the sensitivity of cells to sublethal combinations of cerulenin and rapamycin (Extended Data Fig. 2a,b), indicating that the elevated malonyl-CoA—and not the reduced palmitate—is responsible for this phenotype. Together, these data suggest that TORC1 may be inhibited by malonyl-CoA, in response to perturbations in Acc1 and FAS activity.

Supporting this model, *acc1*^*S1157A*^ cells exhibited significantly reduced TORC1 activity when grown exponentially, and were partially impaired in glutamine-stimulated reactivation of TORC1 in nitrogen-starved cells (Fig. 1e,f). These defects correlated well with significantly increased levels of malonyl-CoA in cells expressing Acc1^S1157A^ (Extended Data Fig. 2c,d). Moreover, combining the *acc1*^*S1157A*^ mutation with an *acc1*^*E392K*^ mutation (also known as *acc1-7-1* ^30^), a temperature-sensitive allele (Extended Data Fig. 2e) that is hypomorphic for malonyl-CoA production at the permissive temperature (Extended Data Fig. 2c,d), suppressed both the elevated malonyl-CoA levels and the reduced TORC1 activity that are observed in *acc1*^*S1157A*^ cells (Extended Data Fig. 2c,d,f,g). Similarly, C-terminal GFP-tagging of Acc1^S1157A^ rendered this allele less active, which significantly reduced the cellular malonyl-CoA levels (Extended Data Fig. 3a,b) and suppressed the cerulenin-sensitivity (Extended Data Fig. 3c), the rapamycin-sensitivity (Extended Data Fig. 3c), and the TORC1 activity defect (Extended Data Fig. 3d,e) that are observed in untagged Acc1^S1157A^-expressing cells. In contrast, C-terminal GFP-tagging of Fas1 lead to cerulenin sensitivity in wild-type cells, and further enhanced this effect in *acc1*^*S1157A*^ cells (Extended Data Fig. 3c).

### FASN inhibition controls mTORC1 activity independently from FA biosynthesis

A similar molecular machinery is responsible for FA biosynthesis in mammalian cells, which differ from yeast by expressing a single fatty acid synthase (FASN) enzyme that catalyses the conversion of malonyl-CoA to palmitate and other FAs (Fig. 1a). Therefore, we sought to investigate if accumulation of malonyl-CoA influences mTORC1 activity also in mammalian cells. To test this, we blocked FASN activity in human HEK293FT cells, either by specific pharmacological inhibition using Fasnall (also known as benzenesulfonate) ^31^ (Fig. 1a,g,h) or cerulenin (Fig. 1a,i,j), or by transient *FASN* knockdown by siRNA transfection (Fig. 1k,l). In accordance with the yeast data, all perturbations suppressed mTORC1 activity, as indicated by decreased phosphorylation of its direct substrates S6K and 4E-BP1 (Fig. 1g-l). As expected, cerulenin or *FASN* knockdown led to a detectable increase in total protein malonylation, detected by immunoblotting with an anti-malonyl-lysine (α-Mal-K) antibody (Fig. 2i,k), indicative of elevated intracellular malonyl-CoA levels ^32,33^. Similar results were obtained by inhibiting or knocking down FASN in MEFs (mouse embryonic fibroblasts) (Extended Data Fig. 4a-c), MCF-7 human breast cancer cells (Extended Data Fig. 4d,e), WI-26 human lung fibroblasts (Extended Data Fig. 4f), or U2OS human osteosarcoma cells (Extended Data Fig. 4g,h), showing that this effect is not cell type-specific. Further supporting that malonyl-CoA is able to inhibit mTORC1, exogenous addition of this metabolite ^34,35^ caused a significant drop in mTORC1 activity in HEK293FT cells (Extended Data Fig. 5a,b). Importantly, FASN inhibition specifically affected mTORC1, but not mTORC2, as treatment with Fasnall or cerulenin did not significantly alter phosphorylation of Akt, a typical mTORC2 substrate in human cells (Extended Data Fig. 5c-f). Similarly, treatment with cerulenin or expression of Acc1^S1157A^ did not inhibit the phosphorylation state of Ypk1^Thr662^, a *bona fide* TORC2 target residue in yeast (Extended Data Fig. 5g-j).

**Figure 2.**
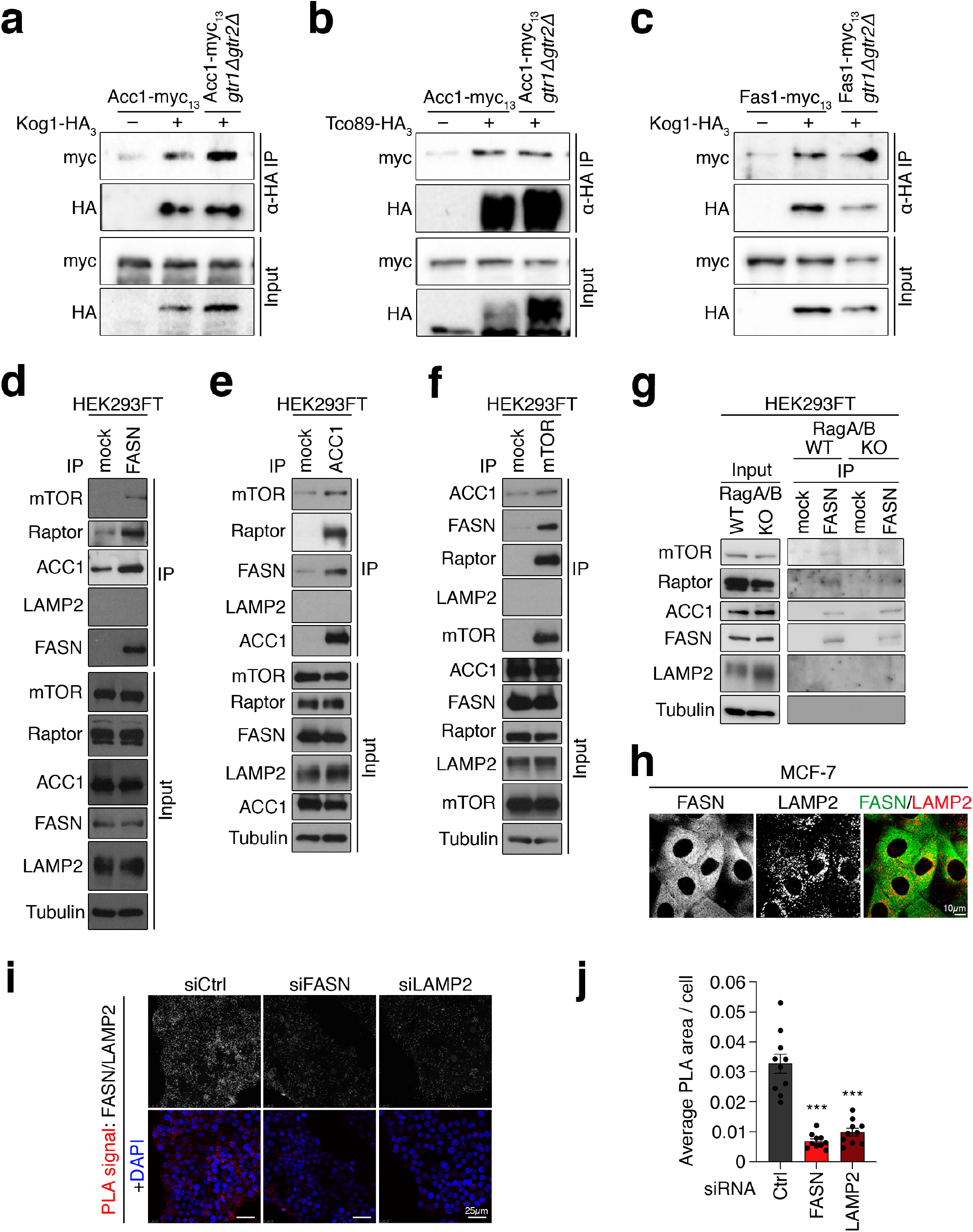
The mTORC1-FASN-ACC1 proteins form reciprocal interactions in yeast and mammalian cells. **(a-b)** Acc1 physically interacts with TORC1 in a Rag-independent manner. Wild-type and *gtr1Δ gtr2Δ* cells expressing genomically tagged Acc1-myc_13_ and untagged (–) or genomically HA_3_-tagged (+) TORC1 subunits Kog1 (a) or Tco89 (b) were grown exponentially. Lysates (Input) and anti-HA immunoprecipitates (α-HA IP) were analysed by immunoblotting with anti-myc and anti-HA antibodies. **(c)** Fas1 physically interacts with TORC1 in a Rag-independent manner. As in (a-b), using wild-type and *gtr1Δ gtr2Δ* cells harbouring genomically tagged Fas1-myc_13_ and untagged (–) or genomically HA_3_-tagged (+) Kog1. **(d)** FASN directly interacts with mTOR, Raptor and ACC1. Endogenous FASN was immunoprecipitated from HEK293FT cell lysates and co-immunoprecipitated proteins were identified by immunoblotting as indicated. **(e)** As in (d), but with ACC1 immunoprecipitation. **(f)** As in (d), but with mTOR immunoprecipitation. **(g)** The interaction between FASN, mTOR/Raptor and ACC1 is independent from the Rags. FASN was immunoprecipitated from WT or RagA/B KO HEK293FT cell lysates and co-immunoprecipitated proteins were identified by immunoblotting as indicated. **(h)** Immunofluorescence of FASN and LAMP2 (lysosomal marker) in MCF-7 cells shows primarily diffuse cytoplasmic FASN localization. Scale bar = 10 μm. **(i-j)** FASN shows proximity to lysosomes. PLA assays in MCF-7 cells using FASN and LAMP2 antibodies. Specificity of the PLA signal (red dots) verified by FASN and LAMP2 knockdowns. Scale bars = 25 μm (i). Quantification of PLA signal intensity in (j). Data shown as mean ± SEM. *** = p < 0.0005.

Palmitate, the main FASN product, was previously suggested to be important for mTORC1 activity in specialized cell types ^36-38^. We therefore tested if FASN inhibition downregulates mTORC1 due to a decrease in palmitate or an increase in malonyl-CoA by modulating the levels of these two metabolites. Notably, neither culturing of cells in charcoal-stripped FBS that is depleted of lipids (Extended Data Fig. 6a,b) nor supplementation of culture media with BSA-conjugated FAs modulated the inhibition of mTORC1 by Fasnall or cerulenin treatment (Fig. 1m and Extended Data Fig. 6b). In contrast, exogenous expression of the constitutively active ACC1^S79A^ mutant demonstrated a synergistic effect with cerulenin towards mTORC1 inhibition, without significantly affecting the response to amino acid starvation (Extended Data Fig. 6c,d), also hinting that the inhibition of mTORC1 is due to elevated malonyl-CoA levels.

A previous study proposed that malonylation of mTOR on lysine 1218 (K1218) negatively impacts mTORC1 activity in endothelial cells ^39^. Therefore, we tested if the accumulation of malonyl-CoA upon FASN/Fas1 inhibition is downregulating mTORC1/TORC1 via this mechanism. However, by using an antibody that detects malonylated lysine (Mal-K) residues on proteins, we were not able to detect malonylation of immunopurified human mTOR or yeast Tor1, from control or cerulenin-treated cells, in our system (Extended Data Fig. 6e,f). In contrast to mTOR, and in agreement with previous proteomics analyses of the human malonylome ^33,40^, lysine malonylation was readily and robustly detectable on immunopurified human FASN and yeast Fas1 from control cells, with the malonylation increasing further in cerulenin-treated cells (Extended Data Fig. 6e,f). Taken together, our data from yeast and mammalian cells indicate that hyperactivation of ACC1 or inhibition of FAS downregulate mTORC1/TORC1 via a mechanism that involves accumulation of malonyl-CoA, independently from mTOR malonylation and from its role in FA biosynthesis.

### Perturbations in yeast and mammalian ACC1/FAS activity control mTORC1 independently of upstream regulatory complexes

To study how malonyl-CoA may be regulating TORC1 activity, we first employed genetic epistasis in yeast. Nutrients such as amino acids regulate TORC1 in part via the Gtr/Rag GTPases ^23^. Expression of constitutively active Acc1^S1157A^, however, rendered not only wild-type cells, but also *gtr1Δ, gtr1*^*S20L*^, *gtr2Δ*, and *gtr2*^*S23L*^ mutant cells cerulenin-sensitive (Extended Data Fig. 2a,b), indicating that Acc1 or its product malonyl-CoA may impinge on TORC1 independently of Gtr1/2. To further corroborate this assumption, we assayed the effects of hyperactive Acc1^S1157A^ expression on TORC1 upon combined loss of Gtr1 and Gtr2, which, in control experiments, did not prevent the Acc1^S1157A^-mediated increase in malonyl-CoA (Extended Data Fig. 7a). As expected, either expression of Acc1^S1157A^ or loss of Gtr1/2 reduced TORC1 activity (Extended Data Fig. 7b,c). Expression of Acc1^S1157A^ in the Gtr1/2 double-mutant background decreased TORC1 activity further, suggesting that Acc1^S1157A^ acts on TORC1 independently of Gtr1/2 (Extended Data Fig. 7b,c). Furthermore, the presence of the Acc1^S1157A^ allele did not affect the vacuolar localization of GFP-tagged Tor1 or Gtr1 (Extended Data Fig. 8a,b), while expression of constitutively active Gtr1^Q65L^ was unable to revert the Acc1^S1157A^-mediated TORC1 inhibition (Extended Data Fig. 7d,e). Together, these results establish that the effects of uncontrolled, Acc1-dependent malonyl-CoA synthesis on TORC1 do not require the presence of the Rag GTPases in yeast cells.

The heterodimeric Rag GTPases and the TSC complex are two major signalling hubs upstream of mTORC1 in mammalian cells ^8,9,41-44^. Therefore, we next tested if malonyl-CoA regulates mTORC1 through one of these complexes. In agreement with our yeast data, FASN inhibition by Fasnall or cerulenin decreased mTORC1 activity to a similar extent in wild-type, RagA/B, or RagC/D double knockout cells, suggesting that it acts independently from the Rags (Extended Data Fig. 7f-h). Furthermore, Fasnall treatment was equally capable of downregulating mTORC1 in both control cells and cells that exogenously express an ‘active’-locked RagA/C mutant dimer, which partially prevents mTORC1 inactivation upon amino acid starvation (Extended Data Fig. 7i,j). Finally, TSC1 knockout cells, that demonstrate a compromised response to amino acid removal ^42^, showed a similar mTORC1 downregulation when treated with Fasnall (Extended Data Fig. 7k,l) or cerulenin (Extended Data Fig. 7m,n), compared to wild-type controls. Because inhibition of FASN can inactivate mTORC1 independently of the two major upstream signalling hubs, these data suggest that malonyl-CoA accumulation may act downstream of these regulatory complexes, possibly by directly acting on mTORC1.

The lysosomal recruitment of mTORC1 by the Rag GTPases is an important aspect of its reactivation in response to AA re-supplementation ^8,41^. Interestingly, we observed that FASN inhibition by Fasnall or cerulenin also led to a significant dissociation of mTOR from lysosomes, as assayed by its colocalization with the lysosomal marker LAMP2 (Extended Data Fig. 8c-f). However, since active RagA/C overexpression or loss of TSC1 (both of which were previously shown to retain lysosomal mTOR even upon starvation ^8,41,42^) still allow downregulation of mTORC1 upon treatment with FASN inhibitors (Extended Data Fig. 7i-n), we conclude that the delocalization of mTOR away from lysosomes is not the cause of its inactivation.

### The mTOR-FASN-ACC1 proteins form reciprocal interactions in yeast and mammalian cells at multiple subcellular locations

Because our data in yeast and mammalian cells indicated that the modulation of the carbon flux through ACC and FAS affected TORC1/mTORC1 in a Rag GTPase-independent manner (Extended Data Fig. 7a-j), we entertained the idea that either or both of these enzymes may directly interact with TORC1/mTORC1. In support of this assumption, we found myc-tagged Acc1 to interact with the yeast TORC1 subunits Kog1-HA or Tco89-HA both in the presence and absence of Gtr1/2 (Fig. 2a,b). Similarly, myc-tagged Fas1 also interacted with Kog1-HA even in the absence of Gtr1/2 (Fig. 2c). Notably, reciprocal interactions between endogenous FASN, ACC1 and mTOR proteins were also detected in mammalian cells (Fig. 2d-f), with mTOR and ACC1 co-immunoprecipitating with FASN even in RagA/B double knockout cells that lack an intact Rag GTPase dimer (Fig. 2g).

In conditions of amino acid sufficiency, the active Rag GTPase complex recruits mTORC1 to the lysosomal surface, while in cells that lack a functional Rag GTPase dimer mTORC1 demonstrates a diffuse cytoplasmic localization pattern ^8,41,42^. Hence, our observation that FASN inhibition downregulates mTORC1 independently from the Rags (Extended Data Fig. 7a-j), and that mTOR interacts with FASN and ACC to the same extent in both Rag-proficient (in which mTOR is lysosomal) and Rag-deficient cells (in which mTOR is non-lysosomal) (Fig. 2g) prompted us to investigate where FASN localizes in cells. In line with the fact that FA biosynthesis takes place primarily in the cytoplasm, and in agreement with publicly available protein localization databases (e.g., the human protein atlas ^45^), endogenous FASN immunostaining showed a diffuse cytoplasmic signal, a portion of which colocalized with the lysosomal marker LAMP2 (Fig. 2h and Extended Data Fig. 8g). Indeed, proximity ligation assay (PLA) experiments using antibodies against endogenous FASN and LAMP2 proteins confirmed that a subpopulation of FASN molecules specifically localize at—or proximal to—the lysosomal surface (Fig. 2i,j). Taken together, these data show that FASN (and possibly also ACC1) physically interact with mTOR both at the lysosomal surface and away from it, suggesting that FASN inhibition and the subsequent accumulation of malonyl-CoA may be directly affecting mTORC1 activity at multiple subcellular locations.

### Molecular dynamics simulations indicate mal-CoA can bind to the mTOR catalytic pocket, similar to ATP

Our genetic, pharmacological, and biochemical data hinted at a possible direct role of malonyl-CoA in the inhibition of mTORC1. Taking into consideration that the adenine-moiety of malonyl-CoA structurally resembles ATP (Fig. 3a), we hypothesized that malonyl-CoA may be inhibiting mTOR by binding to its catalytic pocket. To investigate this possibility further, we looked at the binding of malonyl-CoA, acetyl-CoA, and Coenzyme A (CoA) to mTOR at the atomistic level, by modelling the complexes starting from the crystallographic structure of mTOR bound to ADP (PDB ID: 3JBZ; see methods). Since the interaction between these compounds and mTOR has not been described before, we first hypothesized that the adenine ring, that is present in all compounds, would localise similarly to ADP in the binding pocket of the kinase domain (KD) of mTOR. Thus, the adenine ring of each compound was first aligned to ADP, and then, to find a good arrangement of the lateral chain in the pocket, site-specific docking simulations were performed, allowing the torsion of the lateral chain only (see also methods). For each compound, three best poses were selected considering the most favourable binding energy values (Extended Data Fig. 9a-c), and subsequently used as a starting point to perform all atom Molecular Dynamics (MD) simulations. As expected, ATP remained in the binding pocket during the entire simulation (Fig. 3b,c and Extended Data Video 1). Similarly to ATP, the adenine ring of malonyl-CoA also remained stable in the binding pocket (Fig. 3b,c and Extended Data Video 2). In stark contrast, however, Coenzyme A and acetyl-CoA, quickly (after 15-45 ns and 67-80 ns, respectively) detached from the protein and moved away from the binding pocket of mTOR (Fig. 3c and Extended Data Videos 3, 4). The origin of these differences in interactions between malonyl-CoA and the other CoA-containing compounds with mTOR is likely due to the negatively charged chain (COOH^−^) of its malonyl group that engages in multiple interactions with positively charged mTOR residues just outside the binding pocket (Extended Data Fig. 10a), such as R2168, R2170, and K2187. In contrast, the respective acetyl-CoA (-CH3) and CoA (-SH) groups are not charged and, thus, do not engage in similar interactions (Extended Data Fig. 10b,c). Accordingly, MD simulations and *in silico* mutagenesis of mTOR in the R2168/R2170 residues, that seemingly participate in the stabilization of malonyl-CoA (mTOR^R2168A/R2170A^), hinted at a possible role of these residues in malonyl-CoA binding (Extended Data Fig. 10d and Extended Data Video 5). Intriguingly, these residues are strongly conserved in mTOR over a wide range of organisms, spanning from yeast to humans (Extended Data Fig. 10e).

**Figure 3.**
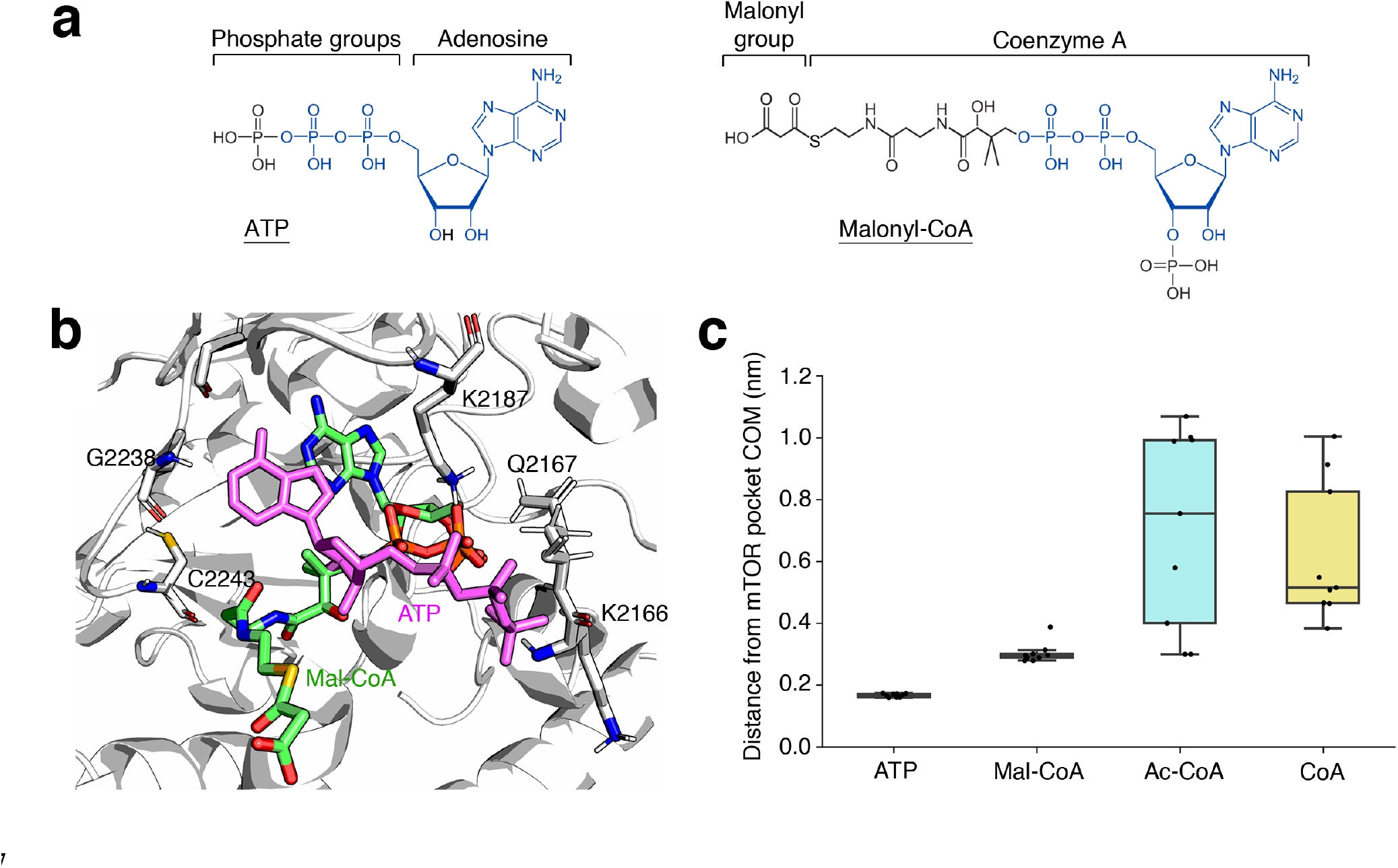
Molecular dynamics simulation of Mal-CoA binding to the mTOR catalytic pocket. **(a)** Chemical structures of ATP (left) and malonyl-CoA (right) highlight structural similarities between the two molecules. Identical parts marked in blue. **(b)** Structural alignment of representative snapshots of malonyl-CoA (green; initial conformation shown) and ATP (magenta) bound to the mTOR catalytic pocket (top view). **(c)** Distances of the ligands from the mTOR binding pocket during the MD simulations. The distances were computed between the center of mass (COM) of the adenine ring and the COM of the amino acid residues defining the pocket. Three replicas were run for each compound, and individual dots represent averages over 100 ns of MD simulation (n = 9). Data in box plots: central line, median; box, interquartile range (IQR) [25^th^ (Q1) - 75^th^ (Q3) percentile]; whiskers, Q3+1.5*IQR and Q1-1.5*IQR.

### Mal-CoA is a direct ATP-competitive inhibitor of mTORC1 activity

To experimentally test our *in silico* analyses, we next sought to investigate whether malonyl-CoA is able to inhibit mTORC1 activity directly, in a cell-free system. Strikingly, classical *in vitro* enzyme kinetics assays, using TORC1 purified from yeast cells and recombinant Lst4 (*i*.*e*. Lst4^Loop 46^) or co-purified Tco89 ^47^ proteins as substrates, showed that this was indeed the case. Similar to what we observed in cells, when added to the *in vitro* kinase (IVK) reaction, malonyl-CoA inhibited TORC1 in a dose-dependent manner with a calculated IC_50_ of 0.73 mM, while the respective IC_50_ values for acetyl-CoA and CoA were 2.98 mM and above our detection limit of 6 mM, respectively (Fig. 4a,b). Accordingly, mTORC1 IVK assays with increasing amounts of malonyl-CoA, acetyl-CoA, and CoA, using immunoprecipitated mTORC1 from mammalian cells and recombinant 4E-BP1 as substrate, revealed a dose-dependent inhibition of mTORC1 by malonyl-CoA (IC_50_ = 0.23 mM), with acetyl-CoA being substantially less potent (IC_50_ = 1.03 mM), and CoA being unable to inhibit mTORC1 activity under our experimental conditions (IC_50_ > 5 mM) (Fig. 4c-e). As a control, malonyl-CoA addition to cell lysates before immunoprecipitation, did not influence mTORC1 stability, as indicated by the interaction of mTOR with RAPTOR and mLST8 proteins (Extended Data Fig. 10f). Overall, our data confirm that malonyl-CoA (and, to a lesser extent, acetyl-CoA) can act as a direct mTORC1 inhibitor, without affecting complex composition.

**Figure 4.**
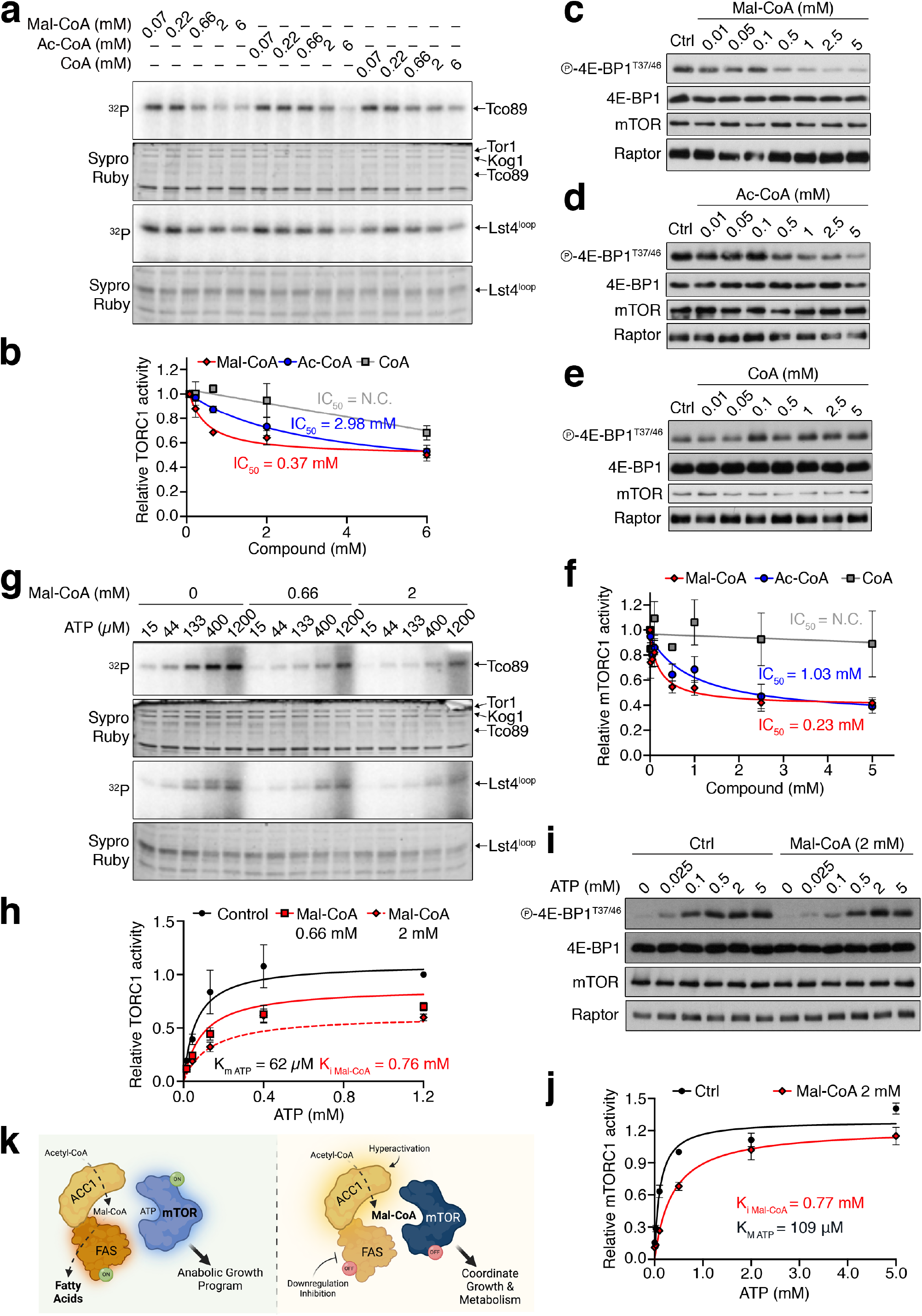
Mal-CoA is a direct ATP-competitive inhibitor of mTORC1. **(a-b)** Malonyl-CoA, and to a lesser extent acetyl-CoA, inhibit TORC1 activity *in vitro*. IVK assays using TORC1 purified from yeast with recombinant Lst4^Loop^ and co-purified Tco89 proteins as substrates, in the presence of increasing Mal-CoA, Ac-CoA, or CoA amounts. Substrate phosphorylation detected by autoradiography (^32^P). Total protein detected by Sypro Ruby staining (a). Quantification of TORC1 activity (Lst4^Loop^ phosphorylation) in (b). Curve fitting and IC_50_ calculations performed with GraphPad Prism. Data shown as mean ± SEM, n = 3. **(c-f)** IVKs as in (a), but with mTORC1 purified from HEK293FT cells and recombinant 4E-BP1 protein used as substrate in the presence of increasing Mal-CoA (c), Ac-CoA (d), or CoA (e) amounts. 4E-BP1 phosphorylation detected by immunoblotting. Quantification of mTORC1 activity in (f). Curve fitting and IC_50_ calculations performed with GraphPad Prism. Data shown as mean ± SEM, n = 4. **(g-h)** Malonyl-CoA inhibits TORC1 in an ATP-competitive manner. IVKs performed as in (a) using increasing ATP concentrations without or with 0.66 or 2 mM Mal-CoA. Quantification of TORC1 activity (Lst4^Loop^ phosphorylation) in (h). Curve fitting and calculations of K_m ATP_ and K_i Mal-CoA_ performed with GraphPad Prism. Data shown as mean ± SEM, n = 3. **(i-j)** IVKs as in (c) using increasing ATP concentrations, in the presence or absence of 2 mM Mal-CoA (i). Quantification of mTORC1 activity in (j). Curve fitting and calculations of K_m ATP_ and K_i Mal-CoA_ performed with GraphPad Prism using a mixed inhibition model. Data shown as mean ± SEM, n = 4. **(k)** Model of mTOR inhibition by Mal-CoA. When the FA biosynthesis machinery is active, ACC1 converts Ac-CoA to Mal-CoA, which is, in turn, rapidly converted to palmitate by FASN (left). In contrast, when ACC1 is hyperactive or FASN is downregulated, accumulating Mal-CoA competes with ATP for binding to proximal mTOR molecules causing its inactivation. Hence, by complexing with ACC1 and FASN, mTORC1 functions as a direct sensor for Mal-CoA to adjust growth and coordinate cellular metabolic activity in response to decreased cellular FA biosynthesis capacity (right).

The resemblance of malonyl-CoA to ATP and our MD simulation experiments suggested that malonyl-CoA may inhibit TORC1/mTORC1 directly, through competition with ATP. In strong support of this hypothesis, IVK assays performed with increasing ATP concentrations, using yeast TORC1 (Fig. 4g) or human mTORC1 complexes (Fig. 4i), when subjected to regression analyses using the GraphPad Prism curve fitting program, indicated that the behaviour of malonyl-CoA matched best with that of an ATP-competitive TORC1/mTORC1 inhibitor with a calculated K_i_ of 0.76 mM or 0.77 mM, for yeast or human complexes, respectively (Fig. 4h,j). In sum, our data reveal that malonyl-CoA is a direct, ATP-competitive inhibitor of mTORC1 in both yeast and human cells, thus serving as a key metabolite that directly connects the cellular FA biosynthetic capacity to the activity of the main cellular metabolic regulator (Fig. 4k).

## Discussion

A key characteristic of mTORC1 is that it forms homeostatic feedback loops, acting both as a molecular sensor and as a regulator of individual biosynthetic processes. For instance, mTORC1 is the master controller of protein synthesis via directly phosphorylating S6K and 4E-BP1, while it also senses the sufficiency of amino acids and energy, thus ensuring that cells only make proteins when all building blocks are available ^1^. Whereas mTORC1 was previously described to regulate lipid biosynthesis at several levels, by controlling the activity and localization of Lipin-1 ^48-50^ and the activity of SREBP transcription factors ^51,52^, how the FA biosynthetic metabolic status signals back to regulate mTORC1 activity is less clear. Here, we report that this interplay also happens in the opposite direction, with key components of the core FA biosynthesis machinery (namely ACC1 and FASN) interacting directly with mTOR, and their activity being tightly connected to that of mTORC1. This way, the FA biosynthesis capacity of a cell is closely coupled to cell growth, metabolism, and other mTORC1 downstream cellular functions. Accordingly, SREBPs are the best described transcription factors controlling FASN expression ^53,54^. By mTORC1 controlling SREBP activity, and thereby FASN levels ^11^, our findings reveal a positive feedback loop between mTORC1 and FASN that could function to sustain lipid biosynthesis when conditions are optimal.

Endogenous metabolites are known to control the activity of key signalling molecules by directly binding to them and modifying their structure and function. For instance, binding of four cAMP molecules to the two regulatory subunits of PKA (protein kinase A) cause their dissociation from the catalytic PKA subunits, which are then activated and directly regulate downstream effectors, such as the CREB transcription factor, to modulate cellular metabolism ^55-57^. Similarly, under low energy conditions, AMP allosterically activates AMPK by binding directly to its γ subunit ^17^. Along the same lines, a recent study showed that inositol directly competes with AMP for binding to AMPKγ, thereby allosterically inhibiting AMPK enzymatic activity, with low inositol driving the AMPK-dependent mitochondrial fission upon energetic stress ^18^. Our work here describes another example of an endogenous metabolite (i.e., malonyl-CoA) that functions as a direct regulator of a central signalling complex (i.e., mTORC1). However, unlike the allosteric modulation of kinase activities described above, malonyl-CoA acts as a direct, ATP-competitive mTORC1 inhibitor, an attribute that stems from the structural similarity between the CoA moiety and ATP. Molecular dynamics simulation experiments also support a model where the charged malonyl group helps to stabilize malonyl-CoA binding to mTOR via interactions with residues just outside its catalytic pocket. Importantly, we find that this is an ancient mechanism that is present already in yeast cells, and is conserved through evolution all the way to humans.

We were, unfortunately, not able to experimentally test the role of the R2105/R2107 residues of Tor1 (R2168/R2170 in human mTOR) in cells, as the respective alanine mutant protein was extremely unstable. Moreover, when expression between WT and mutant Tor1 was artificially matched (by introducing additional DNA/cDNA copies to cells), the latter showed diminished Kog1/Raptor binding in co-immunoprecipitation experiments (data not shown). Therefore, in addition to participating in the stabilization of malonyl-CoA in the mTOR catalytic pocket (as shown from our *in silico* mutagenesis and MD simulation analyses), these residues seemingly also play important roles in mTORC1 formation and mTOR stability.

While the absolute concentration of malonyl-CoA in yeast and mammalian cells has not been accurately measured before, our IVK experiments show that malonyl-CoA inhibits mTORC1 with IC_50_ = 0.23 mM and K_i_ = 0.77 mM (IC_50_ = 0.23 mM, K_i_ = 0.76 mM, for yeast TORC1). Although it is possible that the average intracellular malonyl-CoA concentration does not increase to this extent, the direct physical association between ACC1/FASN and mTOR/Raptor indicates that, upon FASN blockage or ACC1 hyperactivation, a local increase in malonyl-CoA levels could inhibit proximal mTORC1. Because FASN can interact with both lysosomal and non-lysosomal mTOR, perturbations in its activity are able to control all subpopulations of mTORC1 in cells. Such metabolic proximity channelling principles have been described before ^58^ and can facilitate efficient transfer of a metabolite from one enzyme to the next, thus bypassing the need for alterations in total intracellular metabolite levels.

Because of its central role in FA biosynthesis, FASN has emerged as a critical player in cancer cell metabolism, growth and survival ^13,59-61^, with several FASN inhibitors currently being tested in clinical trials ^62^. Interestingly, previous work suggested that the accumulation of malonyl-CoA, rather than the inhibition of FASN itself, is the underlying cause in FASN-inhibitor-induced toxicity in breast cancer cells ^34^. In the present study, we find that FASN inhibition also leads to mTORC1 downregulation due to malonyl-CoA accumulation, in addition to its well-known role in FA biosynthesis. As mTOR activity is commonly dysregulated in the majority of human cancers, our work raises the plausible hypothesis that part of the beneficial effect of FASN inhibition in cancer treatment may be due to the concomitant drop in mTORC1 signalling ^63^. Of note, in our experiments, mTORC1 and FASN inhibitors demonstrated synergistic effects in yeast, even when combined at sublethal doses for each individual compound. These data are in agreement with a previous report showing synthetic lethality of cerulenin and rapamycin in cancer cell lines ^64^. In sum, our findings identify a direct connection between the core FA biosynthesis machinery and mTORC1 activity, reveal novel concepts of how metabolic signalling is coordinated in cells, and provide the basis for the development of advanced therapeutic tools to treat human conditions that are linked to hyperactive mTORC1 signalling.

## Methods

### Yeast culture

Yeast cells were grown in liquid to exponential growth phase in SC medium (1.7 g/l yeast nitrogen base (#1545, CONDA) 5 g/l ammonium sulfate (#4808211, MP Biomedicals), 20 g/l glucose (#1422, AppliChem), 2 g/l amino acid dropout -His (#D9520, US Biological), 35 mg/l histidine (#A1341, AppliChem)) at 30°C, unless otherwise stated in the figure legends. All yeast strains used in this study are listed in Suppl. Table 1.

### Yeast culture treatments

For starvation/re-addition experiments, yeast cells growing in exponential phase were filtered and shifted to prewarmed (30 °C) nitrogen starvation medium (1.7 g/l yeast nitrogen base, 20 g/l glucose) for 20 min. Subsequently, glutamine (#119951000, Acros) was added to a final concentration of 3.3 mM (using a 50x stock solution). Treatment with the pharmacological FAS inhibitor cerulenin (#C2389, Sigma-Aldrich) was carried out by adding the drug directly to the cell cultures at the concentration and times indicated in the figure legends.

### Mammalian cell culture

All cell lines were grown at 37 °C, 5% CO_2_. Human female embryonic kidney HEK293FT cells (#R70007, Invitrogen; RRID: CVCL_6911), human female breast adenocarcinoma MCF-7 cells (#HTB-22, ATCC; RRID:CVCL_0031), immortalized mouse embryonic fibroblasts (MEFs), and human female bone osteosarcoma U2OS cells (#HTB-96, ATCC; RRID:CVCL_0042) were cultured in high-glucose Dulbecco’s Modified Eagle Medium (DMEM) (#41965-039, Gibco), supplemented with 10% FBS (#F7524, Sigma; #S1810, Biowest). MCF-7 cells were also supplemented with 1x NEAA (non-essential amino acids) (#11140-035, Gibco). Human male diploid lung WI-26 SV40 fibroblasts (WI-26 cells; #CCL-95.1, ATCC; RRID: CVCL_2758) were cultured in DMEM/F12 GlutaMAX medium (#31331093, Thermo Fisher Scientific), containing 10% FBS. All media were supplemented with 1x Penicillin-Streptomycin (#15140-122, Gibco).

HEK293FT cells were purchased from Invitrogen. Wild-type control immortalized MEFs were a kind gift of Kun-Liang Guan (described in ^65^). U2OS cells were a kind gift of Nils-Göran Larsson. The identity of the WI-26 cells was validated using the Short Tandem Repeat (STR) profiling service, provided by Multiplexion GmbH. The identity of the HEK293FT and MCF-7 cells was validated by the Multiplex human Cell Line Authentication test (Multiplexion GmbH), which uses a single nucleotide polymorphism (SNP) typing approach, and was performed as described at www.multiplexion.de. All cell lines were regularly tested for *Mycoplasma* contamination, using a PCR-based approach and were confirmed to be *Mycoplasma*-free.

### Mammalian cell culture treatments

Amino acid starvation experiments were carried out as described before ^42^. Treatments with the pharmacological FASN inhibitors Fasnall (#SML1815, Sigma-Aldrich) and cerulenin (#C2389, Sigma-Aldrich) were performed by adding the drugs directly to the media at the concentrations and for times indicated in the figure legends, with DMSO (#4720.1, Roth) used as control. For lipid depletion experiments, cells were cultured in media containing 10% charcoal-stripped FBS (CD-FBS) (#A3382101, Thermo Fisher Scientific), instead of full FBS, for 24h prior to treatments with FASN inhibitors. For fatty acid supplementation, each fatty acid was first conjugated to 10% fatty-acid-free BSA (bovine serum albumin fraction V) (#10735086001, Roche) for 1 hour at 50°C in a 50:50 volumetric ratio. The BSA-conjugated C16:0, C16:1, C18:1, and C18:2 fatty acids (100 μM) were then added to the media both 16 h prior to and at the start of the treatment with FASN inhibitors. Exogenous Mal-CoA treatments were performed by adding 250 μM malonyl-CoA lithium salt (#M4263, Sigma-Aldrich) in the culture media for 30 min before cell lysis.

### Antibodies

All antibodies used in this study are listed in Suppl. Table 2.

### Plasmids and molecular cloning

The pcDNA3-FLAG-hRagA^QL^ (Q66L) and hRagC^SN^ (S75N) vectors expressing the constitutively-active Rag GTPases were described previously ^42^. The pcDNA3-FLAG-Luc control vector was described in ^66^. To generate the pcDNA3-FLAG-ACC1 expression vector, the long ACC1 isoform 4 (Uniprot ID Q13085-4; not used in this study) was PCR amplified from cDNA (prepared from MCF-7 cells) using appropriate primers and cloned in the XhoI/XbaI restriction sites of pcDNA3-FLAG. Next, the canonical, shorter ACC1 isoform 1 (Uniprot ID Q13085-1) was generated using the isoform 4 expression vector as a template and appropriate PCR primers, and cloned in the XhoI/BglII restriction sites of pcDNA3-FLAG-ACC1^Iso4^, thus replacing the long N-terminal part with that of ACC1 isoform 1. The respective pcDNA3-FLAG-ACC1^S79A^ (isoform 1) plasmid was generated by site-directed mutagenesis using appropriate primers, and the insert was cloned in the XhoI/BglII restriction sites of pcDNA3-FLAG-ACC1. The pETM-11-4E-BP1 vector, used to express His_6_-tagged 4E-BP1 in bacteria, was generated by PCR-amplifying human 4E-BP1 from cDNA (prepared from HEK293FT cells) using appropriate primers and cloned in the NcoI-NotI restriction sites of pETM-11. The integrity of all constructs was verified by sequencing. All DNA oligonucleotides used in this study are listed in Suppl. Table 3.

### mRNA isolation and cDNA synthesis

Total mRNA was isolated from cells using a standard TRIzol/chloroform-based method (#15596018, Thermo Fisher Scientific), according to manufacturer’s instructions. For cDNA synthesis, mRNA was transcribed to cDNA using the RevertAid H Minus First Strand cDNA Synthesis Kit (#K1631, Thermo Fisher Scientific) according to manufacturer’s instructions.

### Plasmid DNA transfections

Plasmid DNA transfections in HEK293FT cells were performed using Effectene transfection reagent (#301425, QIAGEN), according to the manufacturer’s instructions.

### Generation of knockout cell lines

The HEK293FT RagA/B, RagC/D and TSC1 knockout cell lines were generated using the pX459-based CRISPR/Cas9 method, as described elsewhere ^67^. The sgRNA expression vectors were generated by cloning appropriate DNA oligonucleotides (Suppl. Table 3) in the BbsI restriction sites of pX459 (#62988, Addgene). An empty pX459 vector was used to generate matching negative control cell lines. In brief, transfected cells were selected with 3 μg/ml puromycin (#A11138-03, Thermo Fisher Scientific) 48 h post-transfection. Single-cell clones were generated by single cell dilution and knock-out clones were validated by immunoblotting.

### Gene silencing experiments

For mTORC1 activity assays, transient knockdown of *FASN* was performed using a pool of 4 siGENOME siRNAs (Horizon Discoveries). An siRNA duplex targeting the *R. reniformis* luciferase gene (RLuc) (#P-002070-01-50, Horizon Discoveries) was used as control. Transfections were performed using 20 nM siRNA and the Lipofectamine RNAiMAX transfection reagent (#13778075, Thermo Fisher Scientific), according to the manufacturer’s instructions. Cells were harvested or fixed 48 h post-transfection and knockdown efficiency was verified by immunoblotting.

### Yeast genomic manipulation

Site-directed mutagenesis in yeast was performed by CRISPR/Cas9 using the method described in ^68^ with minor optimizations. Gene deletions and genomic tagging were performed with a standard high-efficiency transformation protocol using cassettes amplified from various plasmids of the pFA6a PCR toolbox ^69^, or by mating and tetrad dissection. See Suppl. Table 4 for the full list of the plasmids used.

### Yeast cell lysis and immunoblotting

For yeast protein extractions, 10 ml of cell culture were mixed with TCA (trichloroacetic acid) at a final concentration of 6%. After centrifugation, the pellet was washed with cold acetone and dried in a speed-vac. The pellet was resuspended with an amount of lysis buffer (50 mM Tris-HCl pH 7.5, 5 mM EDTA, 6 M urea, 1% SDS) that was proportional to the OD_600nm_ of the original cell culture. Proteins were extracted by disruption in a Precellys machine in the presence of glass beads. Subsequently, a Laemmli-based sample buffer (350 mM Tris-HCl pH 6.8, 30% glycerol, 600 mM DTT, 10% SDS, 0.2 mg/ml bromophenol blue) was mixed (1:1) with whole cell extracts and boiled at 98°C for 5 minutes. The analysis was carried out by SDS-PAGE using antibodies as indicated in the figure legends. Band intensities were quantified using the ImageJ software.

### Mammalian cell lysis and immunoblotting

For standard SDS-PAGE and immunoblotting experiments, cells form a well of a 12-well plate were lysed in 250 μl of ice-cold Triton lysis buffer (50 mM Tris pH 7.5, 1% Triton X-100, 150 mM NaCl, 50 mM NaF, 2 mM Na-vanadate, 0.011 gr/ml beta-glycerophosphate), supplemented with 1x PhosSTOP phosphatase inhibitors (#4906837001, Sigma-Aldrich) and 1x cOmplete protease inhibitors (#11836153001, Roche), for 10 minutes on ice. Samples were clarified by centrifugation (14.000 rpm, 15 min, 4 °C) and supernatants were boiled in 1x SDS sample buffer (5x SDS sample buffer: 350 mM Tris-HCl pH 6.8, 30% glycerol, 600 mM DTT, 12.8% SDS, 0.12% bromophenol blue). Samples were analysed by SDS-PAGE using specific primary antibodies as indicated in the figures. Band intensities were quantified using the ImageJ software.

### Co-immunoprecipitation (co-IP)

For yeast co-immunoprecipitations experiments, cells were collected by filtration and immediately frozen in liquid nitrogen. Subsequently, the pellets were mechanically disrupted in a FastPrep machine in 50 ml tubes containing 5 ml of ice-cold lysis buffer (50 mM Tris-HCl pH 7.5, 150 mM NaCl, 10% glycerol, 0.1% Nonidet P40), supplemented with 1x EDTA-free protease inhibitor cocktail (#11697498001, Roche), 1x PhosSTOP phosphatase inhibitor (#04906837001, Roche) and 4 ml glass beads. Total cell extracts were recovered from beads and cleared by centrifugation. At this stage, samples were taken for input analysis and denatured with a Laemmli-based sample buffer (350 mM Tris-HCl pH 6.8, 30% glycerol, 600 mM DTT, 10% SDS, 0.2 mg/ml bromophenol blue). 10-20 mg of cleared lysates were incubated with anti-HA pre-conjugated magnetic beads (#88837, Pierce) for 2 hours at 4 °C. The beads were then washed 5 times with high salt lysis buffer (50 mM Tris-HCl pH 7.5, 300 mM NaCl, 10% glycerol, 0.1% Nonidet P40) and eventually resuspended in 20 μl lysis buffer and 20 μl 2x Laemmli buffer. Samples were analysed by SDS-PAGE using specific primary antibodies as indicated in the figures.

For mammalian endogenous protein co-immunoprecipitation experiments, cells of a near-confluent 10 cm dish were lysed in CHAPS IP buffer (50 mM Tris pH 7.5, 0.3% CHAPS, 150 mM NaCl, 50 mM NaF, 2 mM Na-vanadate, 0.011 gr/ml beta-glycerophosphate), supplemented with 1x PhosSTOP phosphatase inhibitors and 1x EDTA-free cOmplete protease inhibitors (11873580001, Roche) for 10 minutes on ice. Samples were clarified by centrifugation (14000 rpm, 15 min, 4 °C) and supernatants were subjected to IP by the addition of 3 μl of each antibody, incubation at 4 °C rotating for 3 h, followed by incubation (4 °C, rotating) with 30 μl of pre-washed Protein A agarose bead slurry (#11134515001, Roche) for an additional hour. Beads were then washed four times with CHAPS IP wash buffer (50 mM Tris pH 7.5, 0.3% CHAPS, 150 mM NaCl, 50 mM NaF) and boiled in 1x SDS loading buffer. A portion of the samples was kept aside as inputs, prior to the addition of antibodies. Samples were analysed by SDS-PAGE and the presence of co-immunoprecipitated proteins was detected by immunoblotting with appropriate specific antibodies.

To test if malonyl-CoA influences mTORC1 complex stability/composition, 1 mM malonyl-CoA lithium salt solution was added to the lysates before immunoprecipitation of mTOR, as described above. For mTORC1 *in vitro* kinase assays, endogenous mTOR was immunopurified from 1x 10 cm dish per condition as described above. To test malonylation of FASN and mTOR, endogenous proteins were immunopurified from 1x 10 cm dish per condition as described above, except for using a high-stringency Triton IP lysis buffer (50 mM Tris pH 7.5, 1% Triton X-100, 500 mM NaCl, 50 mM NaF, 2 mM Na-vanadate, 0.011 gr/ml beta-glycerophosphate, 1x PhosSTOP phosphatase inhibitors, 1x EDTA-free cOmplete protease inhibitors) and washing 3x with Triton IP wash buffer (50 mM Tris pH 7.5, 1% Triton X-100, 500 mM NaCl, 50 mM NaF) and 2x with Tris wash buffer (50 mM Tris pH 7.5) to remove interacting proteins. Protein malonylation was assayed by immunoblotting using a Mal-K-specific antibody.

### Production of recombinant His_6_-tagged 4E-BP1 protein in bacteria

Recombinant His_6_-tagged 4E-BP1 protein was produced by transforming *E. coli* BL21 RP electrocompetent bacteria with the pETM-11-4E-BP1 vector described above, according to standard procedures. In brief, protein expression was induced with IPTG (isopropyl-β-D-thiogalactopyranoside) for 3 h at 30 °C, and His_6_-4E-BP1 was purified using Ni-NTA agarose (#1018244, QIAGEN) and eluted with 250 mM imidazole (#A1073, Applichem).

### Yeast TORC1 kinase activity assays

TORC1 was purified from yeast cells and radioactive *in vitro* kinase assays were performed essentially as previously described ^46^. In brief, kinase reactions (total volume 30 μl) were performed with 400 ng purified His_6_-Lst4^Loop^ protein and 60 ng TORC1 in kinase buffer (50 mM HEPES/NaOH pH 7.5, 150 mM NaCl). To test the effect of Mal-CoA (#M4263, Sigma-Aldrich), Ac-CoA (#A2181, Sigma-Aldrich) and CoA (#C3144, Sigma-Aldrich), the kinase reaction was preincubated for 15 min with 2 μl of each compound from a 15x stock solution. Reactions were started by adding 2 μl of ATP mix (62.5 mM MgCl_2_, 4.5 mM ATP, 0.8 µM [γ-^32^P]ATP (#SRP-501, Hartmann Analytic). For kinase assays with different ATP concentrations, reactions were started by adding 2 µl of serial 3-fold dilutions of a more concentrated ATP mix (72 mM ATP, 0.8 µM [γ-^32^P]ATP), always containing 62.5 mM MgCl_2_. All reactions were carried out at 30°C for 10 min and stopped by the addition of 3x sample buffer (50 mM Tris-HCl pH 6.8, 5% SDS, 0.05% bromophenol blue, 630 mM DTT, 30% glycerol) and heating at 65°C for 10 min. Proteins were separated by SDS-PAGE and stained in-gel with SYPRO Ruby (#S4942, Sigma-Aldrich) as loading control. Substrate phosphorylation was analysed by autoradiography using a Typhoon FLA 9500 phosphorimager (GE Healthcare) and the raw density of the signals was quantified using the gel analysis tool of ImageJ.

### Mammalian mTORC1 kinase activity assays

*In vitro* mTORC1 kinase assays were developed based on previous reports ^70,71^. In brief, endogenous mTORC1 complexes were purified from HEK293FT cells, essentially as described in the ’Co-immunoprecipitation’ section, with minor modifications. Following the last wash step with CHAPS IP wash buffer, beads were washed once with kinase wash buffer (25 mM HEPES pH 7.4, 20 mM KCl) and excess liquid was removed with a Hamilton syringe, with final bead volume being approximately 10 μl. Kinase reactions were prepared by adding 10 μl 3x kinase assay buffer (75 mM HEPES/KOH pH 7.4, 60 mM KCl, 30 mM MgCl_2_) to the beads. To test the effect of Mal-CoA (#M4263, Sigma-Aldrich), Ac-CoA (#A2181, Sigma-Aldrich), and CoA (#C3144, Sigma-Aldrich) on mTORC1 activity, 1 μl of each compound was pre-incubated with the kinase complex for 5 minutes prior to initiation of the reaction. Compound concentrations are indicated in the figures. Reactions were started by adding 10 μl of kinase assay start buffer (25 mM HEPES/KOH pH 7.4, 140 mM KCl, 10 mM MgCl_2_), supplemented with 500 μM ATP (final concentration in the reaction) and 35 ng recombinant His_6_-4E-BP1 substrate. For competition assays, ATP concentrations are described in the respective figures. Reactions were incubated at 30 °C for 30 min, and stopped by the addition of one volume 2x SDS loading buffer and boiling (5 min, 95 °C). Samples were run in SDS-PAGE and the mTORC1-mediated phosphorylation on 4E-BP1^T37/46^ was detected by immunoblotting with a specific antibody (#9459, Cell Signaling Technology). Signals were quantified using the gel analysis tool of ImageJ and shown as phospho-4E-BP1^T37/46^/4E-BP1 ratio.

### Immunofluorescence and confocal microscopy

Immunofluorescence / confocal microscopy experiments were performed as described previously ^42,72^. In brief, cells were seeded on glass coverslips (coated with fibronectin for experiments with HEK293FT cells), treated as described in the figure legends, and fixed with 4% paraformaldehyde (PFA) in 1x PBS (10 min, RT), followed by two permeabilization/washing steps with PBT (1x PBS, 0.1 % Tween-20). Cells were blocked in BBT (1x PBS, 0.1% Tween-20, 1% BSA) for 45 minutes. Staining with anti-mTOR (#2983, Cell Signaling Technology) and anti-LAMP2 (#H4B4, Developmental Studies Hybridoma Bank) primary antibodies diluted 1:200 in BBT solution was performed for 2h followed by three washes with PBT. Next, cells were stained with highly cross-adsorbed fluorescent secondary antibodies (Donkey anti-rabbit FITC, Donkey anti-mouse TRITC; both from Jackson ImmunoResearch) diluted 1:200 in BBT for 1 h. Nuclei were stained with DAPI (#A1001, VWR) (1:1000 in PBT) for 5 min and coverslips were washed three times with PBT solution before mounting on glass slides with Fluoromount-G (#00-4958-02, Invitrogen). All images were captured on an SP8 Leica confocal microscope (TCS SP8 X or TCS SP8 DLS, Leica Microsystems) using a 40x oil objective lens. Image acquisition was performed using the LAS X software (Leica Microsystems).

### Quantification of colocalization

Colocalization analysis in confocal microscopy experiments was performed as in ^43,72^, using the Coloc2 plugin of the Fiji software ^73^. A minimum of 12 representative images captured from 3-4 independent experiments were used per condition and Manders’ colocalization coefficient (MCC) with automatic Costes thresholding ^74-76^ was calculated in individual cells. The area corresponding to the cell nucleus was excluded from the cell ROI to prevent false-positive colocalization due to automatic signal adjustments. MCC is defined as a part of the signal of interest (mTOR), which overlaps with a second signal (LAMP2). Values are displayed as mean ± SEM, and significance was calculated using Student’s t-test (for pairwise comparisons) and one-way ANOVA with *post hoc* Holm-Sidak comparisons using GraphPad Prism.

### Proximity Ligation Assays (PLA)

Proximity of endogenous FASN to lysosomes (LAMP2 as lysosomal marker) was assessed in MCF-7 cells by PLA assays, using the Duolink In Situ Red Starter Kit Mouse/Rabbit (#DUO92101, Sigma-Aldrich), according to the manufacturer’s instructions with minor modifications. To test specificity of the FASN/LAMP2 PLA signal in the respective assays, transient knockdown of FASN or LAMP2 was performed using a reverse transfection protocol with appropriate siRNAs (siGENOME, Horizon Discoveries) and Lipofectamine RNAiMAX transfection reagent, according to the manufacturer’s instructions. Cells were trypsinized 48 h post-transfection, re-seeded in 16-well chamber slides (#171080, Nunc Lab-Tek), and assayed approximately 24 h later. In brief, cells were fixed in 4% PFA (in 1x PBS), washed/permeabilized with PBT (1x PBS, 0.1 % Tween-20), and blocked with Blocking solution from the Duolink kit. Samples were incubated overnight at 4 °C with specific anti-FASN (#PA5-22061, Thermo Fisher Scientific; dilution 1:400) and anti-LAMP2 (#H4B4, Developmental Studies Hybridoma Bank; dilution 1:200) primary antibodies, processed according to the kit instructions and mounted on slides with a drop of DAPI-containing Duolink in situ mounting medium. Images were captured on a Leica TCS SP8 confocal microscope. A minimum of 10 randomly chosen fields were acquired per condition as z-stacks, and total PLA signal was calculated on the maximal projections of the single PLA channel using ImageJ. Data in graphs presented as the average PLA area per cell, with n_siCtrl_ = 1100, n_siFASN_ = 1272, and n_siLAMP2_ = 1311 individual cells. Statistical analysis was performed with GraphPad Prism.

### Protein modelling

From the crystallographic structure of human mTOR (PDB ID: 3JBZ), the missing residues were built by means of the Modeller version 10.0 ^77^, using the amino acid sequences provided by the Uniprot database (UniProtKB: P42345). Next, the kinase domain (KD) region (residues 684-1058) was extracted and used for the docking calculation. The mTOR R2168A/R2170A double mutant was modelled with UCSF Chimera, using the Dynameomics rotamer library ^78^, starting from the mTOR structure described above.

### Ligand modelling

Starting from the x-ray structure of human mTOR in a complex with ADP (PDB ID: 3JBZ) initial placement of the different ligands inside the binding pocket was performed. For ATP, the missing phosphate group to obtain the ATP molecule from ADP was rebuilt manually. For malonyl-CoA, its structure was extracted from PDB ID: 5MY0 and subsequently aligned to the ATP structure in the mTOR catalytic pocket. Similar procedures were used for acetyl-CoA and Coenzyme A, and their coordinates were retrieved from PDB ID: 1MZJ and 4l8A, respectively. All compounds were parameterized using the Ligand Reader & Modeller tool of the CHARMM-GUI software ^79-81^.

### Docking simulations

For each compound, ten docking simulations were performed using the AutoDock software (version 4.2) ^82^. Polar hydrogens and Kollman charges were added to the macromolecule ^83^. Gasteiger charges were added to the ligands ^84^ and, to confine the adenosine group in an ATP-like orientation, all rotatable bonds were blocked except for the lateral chain. The grid dimension was adjusted to 54 × 40 × 40 points, and the ligand-macromolecule interaction maps were computed using AutoGrid ^85^. The automated docking software AutoDock Vina ^86^ was used to calculate the binding affinity of ligands and the mTOR kinase domain. Docking energies were evaluated by using empirical-free energy functions and Lamarckian genetic algorithms ^87^. A regular precision and a rigid ligand-docking were set for each docking run. To assess the stability of each docked pose the energy values obtained by the docking were considered.

### Molecular Dynamics (MD) simulations

For ATP, MD simulations of the human mTOR kinase domain (KD) region (residues 684-1058) (PDB ID: 3JBZ) were started after manual addition of the missing phosphate group to the structure of ADP in complex with two Mg^2+^ ions. For the other compounds (malonyl-CoA, acetyl-CoA, and CoA), the three most favourable poses from the docking calculation were selected as the starting point for the MD simulations. Each system was solvated with the TIP3P water model ^88^ and neutralized with Na^+^ and Cl^−^ ions at physiological concentration (0.15 M). An energy minimization step was performed using the steepest descent algorithm. After the minimization, a NVT equilibration of 50 ps at 300K was performed, using the V-rescale thermostat with a τT = 1 ps and an integration time step of 2 fs ^89^. Then, NPT simulation was run with a time step of 2 fs, using the Parrinello-Rahman barostat ^90^, isotropic coupling and τp = 2 ps. The temperature was kept constant at 310 K. The electrostatic interactions were calculated using the particle mesh Ewald method ^91^ with a cut-off of 1.2 nm. The same cut-off was also applied for the van der Waals interactions. The simulations were performed with GROMACS (version 2020) ^92^ and using the CHARMM-36 force field ^93^. For each complex, we performed three replicate measurements lasting 400 ns, resulting in an aggregated time of 1.2 μs per system.

### Statistical analysis

Statistical analysis and presentation of quantification data was performed using GraphPad Prism (version 9.1.0). Data in graphs shown as mean ± SEM. Box plots: central line, median; box, interquartile range (IQR) [25^th^ (Q1)-75^th^ (Q3) percentile]; whiskers, Q3+1.5*IQR and Q1-1.5*IQR. Significance was calculated using Student’s t-test (for pairwise comparisons) or one-way ANOVA with *post hoc* Holm-Sidak test (pairwise comparisons to controls). Sample sizes (n) and significance values are indicated in figure legends (* p < 0.05, ** p < 0.005, *** p < 0.0005, **** p < 0.00005, ns = non-significant).

## Acknowledgements

We thank Filippo Artoni (MPI-AGE), Malika Jaquenoud, Marie-Pierre Péli-Gulli, and Susanne Stumpe (University of Fribourg) for technical support; Jin Hou (Shandong University) for the generous gift of plasmids; Robbie Loewith (University of Geneva) for providing antibodies; Andrea Annibal (MPI-AGE) for help with preparation of BSA-conjugated lipids; the MPI-AGE FACS & Imaging Core Facility for support with confocal microscopy; and Patrick Giavalisco from the MPI-AGE Metabolomics core for suggestions and support. L.B. acknowledges support from the Boost! program of the Max Planck Society. C.D. is funded by the European Research Council (ERC) under the European Union’s Horizon 2020 research and innovation programme (grant agreement No 757729) and by the Max Planck Society. C.D.V is funded by the Canton of Fribourg and the Swiss National Science Foundation (310030_166474/184671). S.V. acknowledges support from the Swiss National Science Foundation (PP00P3_194807, 189996). This work was supported by grants from the Swiss National Supercomputing Centre under project ID s1030 and s1132.

## Author Contributions

Experimental work: R.N., L.B., J.N., G.F., S.K., P.G., J.R., S.A.F., A.L.; data analysis: R.N., L.B., J.N., G.F., J.R., A.A.T., C.D.V., C.D.; docking and molecular dynamics simulations: J.A., S.V.; project design & conceptualisation: R.N., L.B., S.V., A.A.T., C.D.V., C.D.; project supervision: S.V., A.A.T., C.D.V., C.D.; funding acquisition: S.V., A.A.T., C.D.V., C.D.; figure preparation: R.N., L.B., J.A., C.D.; manuscript draft: C.D.V., C.D., with contributions from all authors. All authors approved the final version of the manuscript and agree on the content and conclusions.

## Declaration of Interests

The authors declare no competing interests.

## Data availability

The data that support the findings of this study (uncropped immunoblots, microscopy pictures, molecular dynamics) are available from the corresponding authors upon reasonable request.

## Code availability

No code was generated in this study.

## Additional Information

Supplementary Information (Extended Data Figures 1-10, Extended Data Videos 1-5, and Supplementary Tables 1-4) is available for this paper.

## Notes

### Competing Interest Statement

The authors have declared no competing interest.

