## Supplementary material for "Malonyl-CoA is an ancient physiological ATP-competitive mTORC1 inhibitor": Suppl Figures 1-10

### Supplementary Information

Supplementary Tables 1-4, Extended Data Videos 1-5 and Extended Data Figures 1-10

#### Supplemental Tables

Suppl. Table 1. Yeast strains used in this study.

Suppl. Table 2. List of antibodies used in this study.

Suppl. Table 3. DNA oligonucleotides used in this study.

Suppl. Table 4. List of plasmids used for yeast experiments in this study.

#### Extended Data Videos

Extended Data Video 1. Molecular dynamics simulation of ATP binding to mTOR.

Extended Data Video 2. Molecular dynamics simulation of malonyl-CoA binding to mTOR.

Extended Data Video 3. Molecular dynamics simulation of CoA binding to mTOR.

Extended Data Video 4. Molecular dynamics simulation of acetyl-CoA binding to mTOR.

Extended Data Video 5. Molecular dynamics simulation of malonyl-CoA binding to $mTOR^{R2168/2170A}$.

#### Extended Data Figures

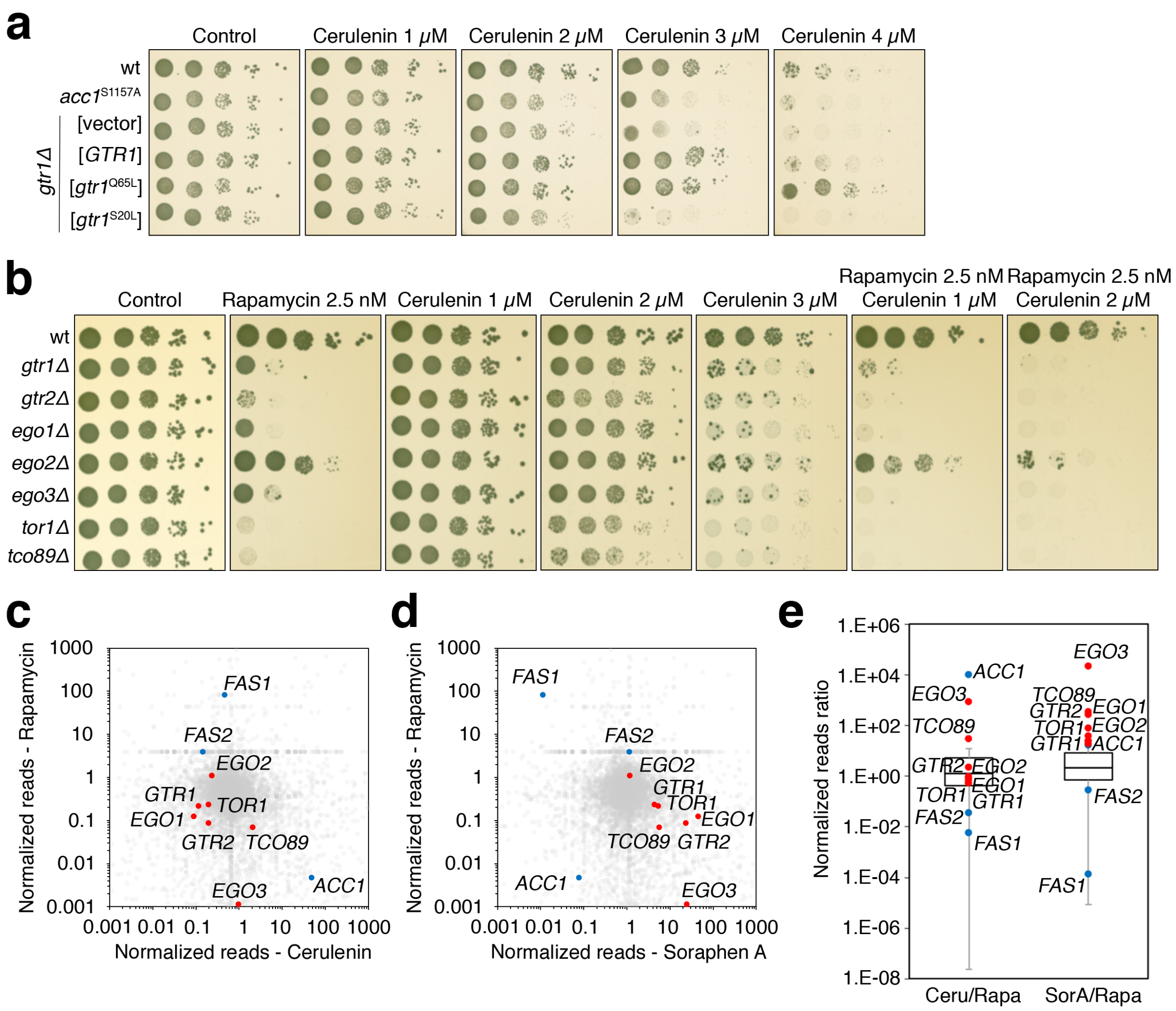


**Extended Data Figure 1. Functional pharmacogenetic interactions between TOR pathway genes and Acc1/Fas1 activity in yeast.**

**(a)** Yeast Rag GTPase mutants that impair or promote TORC1 activity are cerulenin-sensitive or -resistant, respectively. Wild-type (wt), hyperactive $Acc1^{S1157A}$-expressing (*$acc1^{S1157A}$*), and *gtr1Δ* cells that express plasmid-encoded wild-type *GTR1*, *$Gtr1^{Q65L}$* (TORC1-activating), *$Gtr1^{S20L}$* (TORC1-inactivating), or containing an empty vector, were 10-fold serially diluted, spotted on plates with the indicated concentrations of cerulenin or vehicle (Control), and grown at 30°C for 2 days.

**(b)** TORC1 and EGOC mutants are sensitive to cerulenin and hypersensitive to combined cerulenin and rapamycin treatments. Drop spot assays as in (a) using the indicated wt and mutant strains, with plates containing the indicated concentrations of rapamycin and/or cerulenin.

**(c)** Positive correlation between rapamycin- and cerulenin-induced SATAY transposition profiles in TORC1 and EGOC genes. The dot plot shows the ratio of transposition events (reads) per coding gene in rapamycin- and cerulenin-treated versus untreated SATAY libraries previously published in [24,25].

**(d)** Negative correlation between rapamycin- and soraphen-A-induced SATAY transposition profiles in TORC1 and EGOC genes. The dot plot shows the ratio of transposition events (reads) per coding gene in rapamycin- and soraphen-A-treated versus untreated SATAY libraries previously published in [24,25].

**(e)** Box plot summary of the pairwise correlations of the transposon profiles of TORC1 and EGOC genes shown in (c-d).

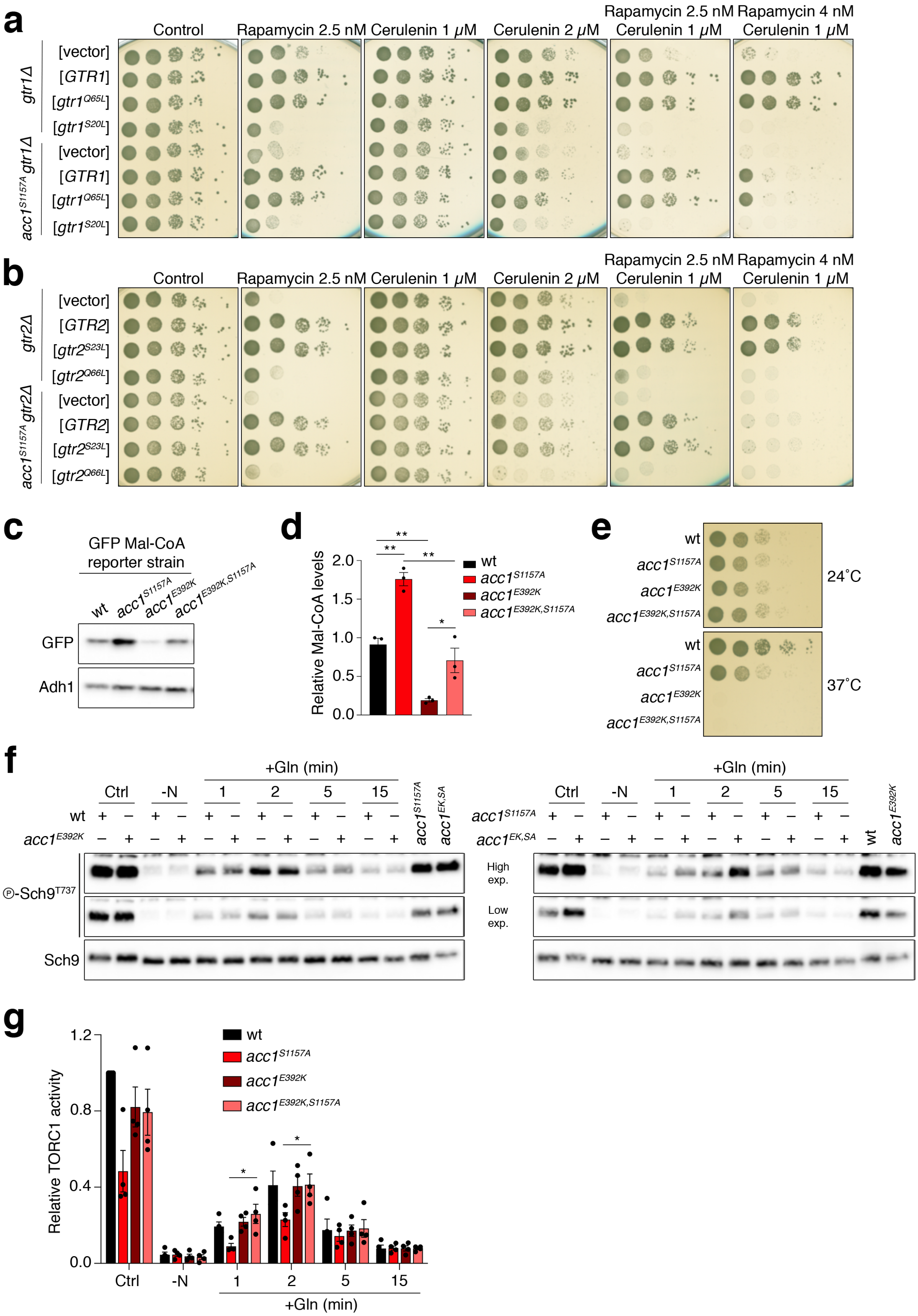
a
Control
Rapamycin 2.5 nM
Cerulenin 1 μM
Cerulenin 2 μM
Rapamycin 2.5 nM Cerulenin 1 μM
Rapamycin 4 nM Cerulenin 1 μM
gtr1Δ
acc1S1157A gtr1Δ
[vector]
[GTR1]
[gtr1Q65L]
[gtr1S20L]
b
gtr2Δ
acc1S1157A gtr2Δ
[GTR2]
[gtr2S23L]
[gtr2Q66L]
c
GFP Mal-CoA reporter strain
wt
acc1S1157A
acc1E392K
acc1E392K,S1157A
GFP
Adh1
d
Relative Mal-CoA levels
e
24°C
37°C
f
+Gln (min)
Ctrl
-N
acc1EK,SA
℗-Sch9T737
Sch9
High exp.
Low exp.
g
Relative TORC1 activity

**Extended Data Figure 2. Perturbations in Acc1 influence Mal-CoA levels and TORC1 activity in yeast cells.**

**(a-b)** Pharmacogenetic interactions of rapamycin and cerulenin indicate a potential inhibitory role of Mal-CoA in TORC1 signalling. *gtr1Δ* and $acc1^{S1157A}$*gtr1Δ* mutant strains containing an empty vector or expressing plasmid-encoded wild-type *GTR1*, $gtr1^{Q65L}$, or $gtr1^{S20L}$ were spotted and grown on plates with the indicated concentrations of rapamycin and/or cerulenin (a). The respective *gtr2Δ* and $acc1^{S1157A}$*gtr2Δ* mutant strains expressing plasmid-encoded wild-type *GTR2*, $gtr2^{Q66L}$, or $gtr2^{S23L}$ are shown in (b).

**(c-d)** Mal-CoA levels in wt and hyperactive $acc1^{S1157A}$ cells are significantly reduced by introduction of the hypomorphic $acc1^{E392K}$ mutation. Mal-CoA levels were assessed by GFP blots in cells grown at 24°C (c) and quantified (d) as in Fig. 1b and Fig. 1c, respectively, n = 3.

**(e)** The $acc1^{E392K}$ mutation causes temperature-sensitive growth even when combined with the $acc1^{S1157A}$ mutation. Exponentially growing cells with the indicated genotypes were spotted on plates (10-fold serial dilutions) and grown for 3 days at 24°C or 37°C.

**(f-g)** Mutation of E392K in $acc1^{S1157A}$ suppresses the TORC1-inhibitory effect of $acc1^{S1157A}$. Immunoblots (f) and quantifications of TORC1 activity (p-Sch9$^{Thr737}$/Sch9) (g) were carried out as in Fig. 1e and Fig. 1f, respectively, n = 4.

Data in graphs shown as mean ± SEM. * = $p < 0.05$, ** = $p < 0.005$

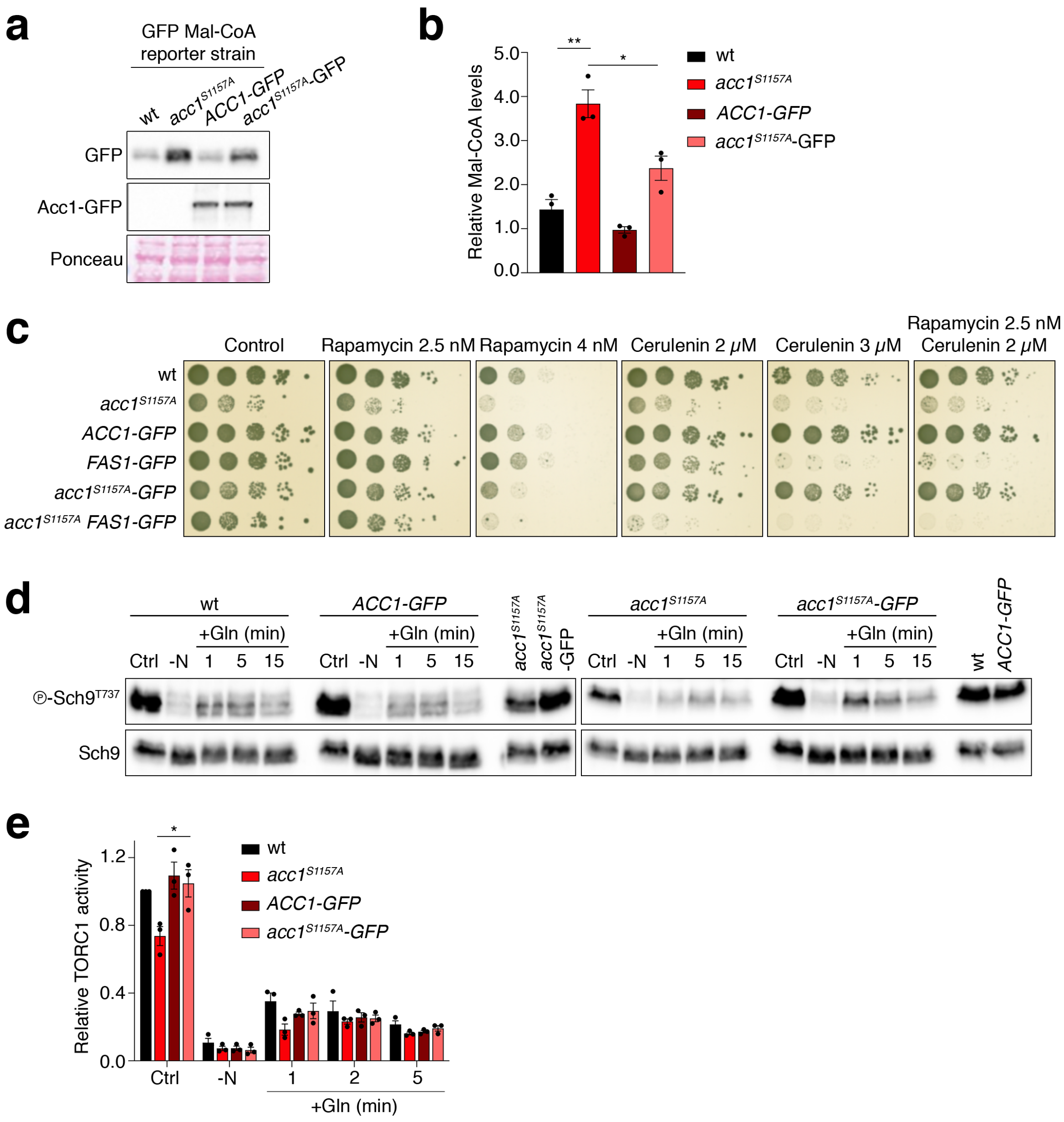


**Extended Data Figure 3. C-terminal tagging of Acc1$^{S1157A}$ restrains its capacity to stimulate Mal-CoA levels and inhibit TORC1 in yeast.**

**(a-b)** C-terminal GFP-tagging of endogenous Acc1$^{S1157A}$ significantly reduces Mal-CoA levels. See Fig. 1b and Fig. 1c for details. Ponceau staining as loading control (a). Quantification of relative Mal-CoA levels (GFP/Ponceau signal) in (b), n = 3.

**(c)** C-terminal GFP-tagging of endogenous Acc1$^{S1157A}$ partially reverts its ability to render cells rapamycin- and cerulenin-sensitive. In control experiments, C-terminal GFP-tagging of endogenous Fas1 rendered wt cells cerulenin-sensitive and further enhanced the cerulenin-sensitivity of *acc1$^{S1157A}$* cells, while marginally affecting the rapamycin-sensitivity of the respective cells. Spot assays with the indicated yeast genotypes and rapamycin and/or cerulenin concentrations performed as in Extended Data Fig. 1a.

**(d-e)** C-terminal GFP-tagging of endogenous Acc1$^{S1157A}$ suppresses its ability to inhibit TORC1 in exponentially growing and Gln-restimulated cells. TORC1 activities assessed as in Fig. 1e (d). Quantification of relative TORC1 activity (p-Sch9$^{Thr737}$/Sch9) in (e), n = 3.

Data in graphs shown as mean ± SEM. * = $p < 0.05$, ** = $p < 0.005$.

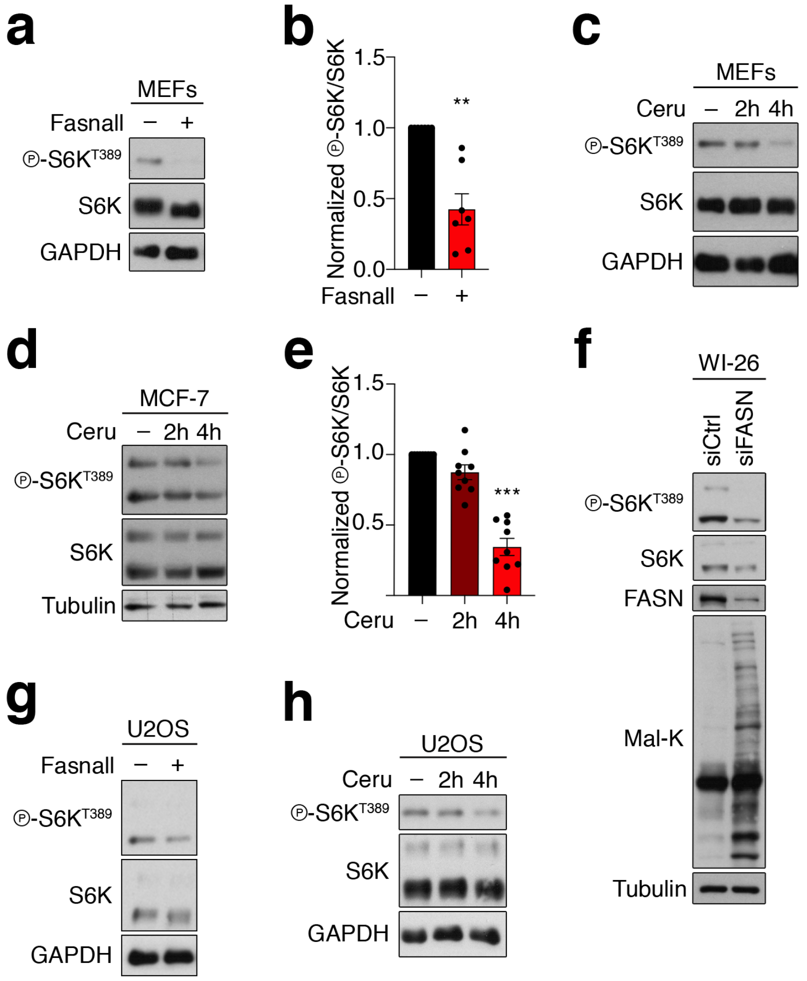


**Extended Data Figure 4. FASN inhibition downregulates mTORC1 in a variety of different mammalian cell lines.**

**(a-b)** Fasnall treatment downregulates mTORC1 activity in MEFs. Cells were treated for 30 min with 25 µM Fasnall or DMSO (–) as control (a). Quantification of mTORC1 activity (p-S6K$^{T389}$/S6K) in (b), n = 7.

**(c)** Cerulenin treatment downregulates mTORC1 activity in MEFs. Cells were treated for 2 or 4 h with 50 µM Cerulenin (Ceru) or DMSO (4 h) as control (–), n = 3.

**(d-e)** Cerulenin treatment downregulates mTORC1 activity in MCF-7 cells. Cells were treated for 2 or 4 h with 50 µM Cerulenin (Ceru) or with DMSO (4 h) as control (–) (d). Quantification of mTORC1 activity (p-S6K$^{T389}$/S6K) in (e), n = 9.

**(f)** *FASN* knockdown downregulates mTORC1 activity and increases Mal-CoA levels in WI-26 fibroblasts. mTORC1 activity assayed by phosphorylation of S6K. Mal-K blots show total protein malonylation, indicative of intracellular Mal-CoA levels.

**(g)** Fasnall treatment downregulates mTORC1 activity in U2OS cells. Cells were treated for 30 or 60 min with 25 μM Fasnall or DMSO as control (–), n = 3.

**(h)** Cerulenin treatment downregulates mTORC1 activity in U2OS cells. Cells were treated for 2 or 4 h with 50 μM Cerulenin (Ceru) or DMSO (4 h) as control (–).

Data in graphs shown as mean ± SEM. ** = $p < 0.005$, *** = $p < 0.0005$.

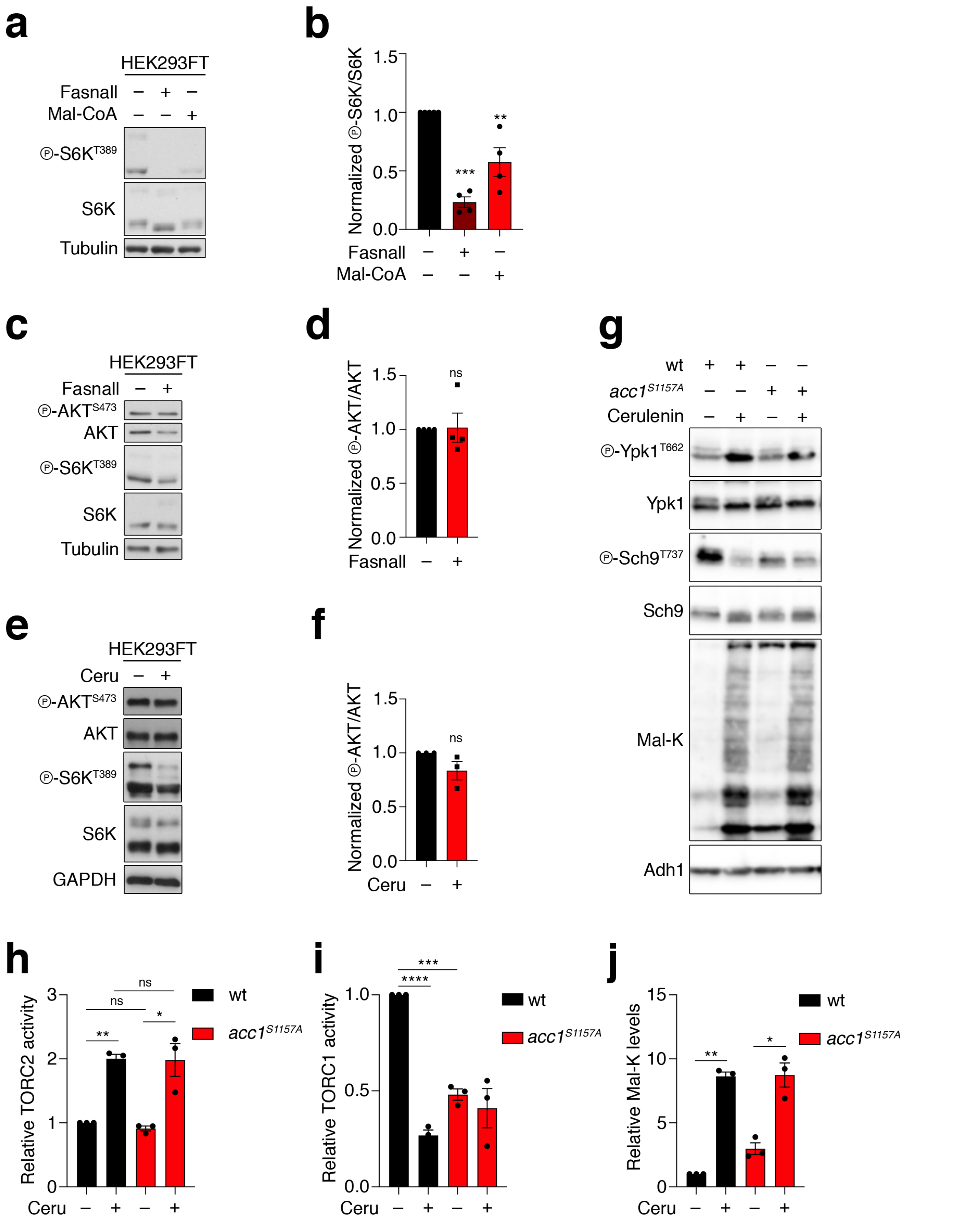


**Extended Data Figure 5. Exogenous Mal-CoA or perturbations in FASN/Fas1 or yeast Acc1 activity specifically downregulate mTORC1, but not mTORC2.**

**(a-b)** Exogenous Mal-CoA addition is capable of downregulating mTORC1. HEK293FT cells were treated with 25 μM Fasnall or 250 μM Mal-CoA for 30 min. mTORC1 activity was monitored by S6K phosphorylation (a). Quantification of mTORC1 activity (p-S6K$^{S389}$/S6K), in (b), n = 4.

**(c-d)** Fasnall treatment does not influence mTORC2 activity in HEK293FT cells, assessed by immunoblotting for AKT phosphorylation. Cells were treated for 30 min with 25 µM Fasnall or DMSO (–) as control (c). Quantification of mTORC2 activity (p-AKT$^{S473}$/AKT) in (d), n = 4.

**(e-f)** Cerulenin treatment does not influence mTORC2 activity in HEK293FT cells, assessed by immunoblotting for AKT phosphorylation. Cells were treated for 4 h with 50 µM Cerulenin (Ceru) or DMSO (–) as control (e). Quantification of mTORC2 activity (p-AKT$^{S473}$/AKT) in (f), n = 3.

**(g-j)** Expression of the *acc1$^{S1157A}$* allele or treatment of yeast cells with cerulenin (20 µM, 2 h) does not downregulate TORC2 activity. Immunoblotting with the indicated antibodies using lysates from control (–) or cerulenin-treated (+) wild-type (wt) or *acc1$^{S1157A}$* mutant cells. Phosphorylation of Ypk1 used as TORC2 read-out. Mal-K blots show total protein malonylation, indicative of intracellular Mal-CoA levels (g). Quantification of TORC2 activity (p-Ypk1$^{T662}$/Ypk1), TORC1 activity (p-Sch9$^{T737}$/Sch9), and lysine malonylation (Mal-K/Adh1) in (h), (i), (j), respectively, n = 3

Data in graphs shown as mean ± SEM. * = $p < 0.05$, ** = $p < 0.005$, *** = $p < 0.0005$, **** = $p < 0.00005$.

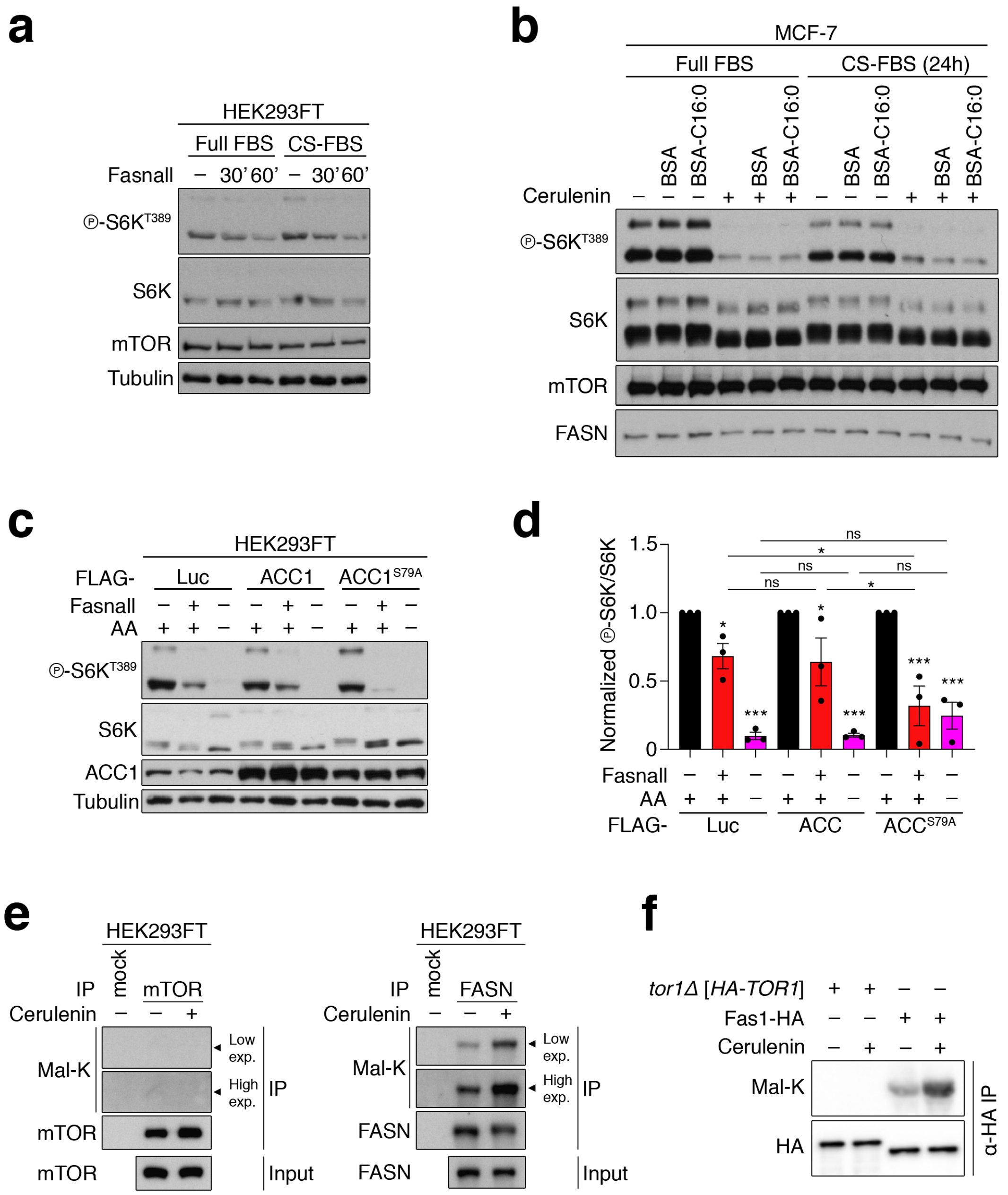

**Extended Data Figure 6. FASN inhibition downregulates mTORC1 activity independently from lipid availability and mTOR/Tor1 malonylation.**

**(a)** Depletion of exogenous lipid sources does not influence the mTORC1 response to FASN inhibition. HEK293FT cells were cultured in full FBS- or charcoal-stripped FBS (CS-FBS)-containing media for 24 h, and then treated with 25 µM Fasnall for 30 or 60 min, or DMSO as control (–). mTORC1 activity assessed by phosphorylation of S6K.

**(b)** Immunoblots with lysates from control (–) or Cerulenin-treated (50 µM, 4 h) MCF-7 cells, supplemented with BSA-conjugated palmitate (C16:0) or BSA as control. Cells were cultured in full

FBS- or charcoal-stripped FBS (CS-FBS)-containing media for 24 h prior to treatments. mTORC1 activity assayed by phosphorylation of S6K.

**(c-d)** Exogenous expression of a hyperactive ACC1 mutant ($ACC1^{S79A}$) cooperates with FASN inhibition to downregulate mTORC1 in HEK293FT cells, without influencing the response to AA starvation. Cells were transfected with vectors expressing FLAG-tagged WT or S79A ACC1, or Luciferase (Luc) as control, and treated with Fasnall (25 μM, 30 min) or AA-starvation media (1 h) as indicated. mTORC1 activity assayed by phosphorylation of S6K (c). Quantification of mTORC1 activity ($p\text{-}S6K^{T389}/S6K$) in (d). Data shown as mean ± SEM, n = 3. * = $p < 0.05$, *** = $p < 0.0005$.

**(e)** No detectable mTOR malonylation in HEK293FT cells. Endogenous mTOR (left) or FASN (right) proteins were immunopurified from control or cerulenin-treated (50 μM, 4 h) cells, and protein malonylation was assayed using an anti-malonyl-lysine (Mal-K) antibody.

**(f)** Tor1 is not malonylated in yeast cells. N-terminally HA-tagged Tor1 or C-terminally HA-tagged Fas1 were immunopurified from control (–) or cerulenin-treated (10 μM, 2 h) cells grown in exponential phase. Protein malonylation was assessed using an anti-malonyl-lysine (Mal-K) antibody.

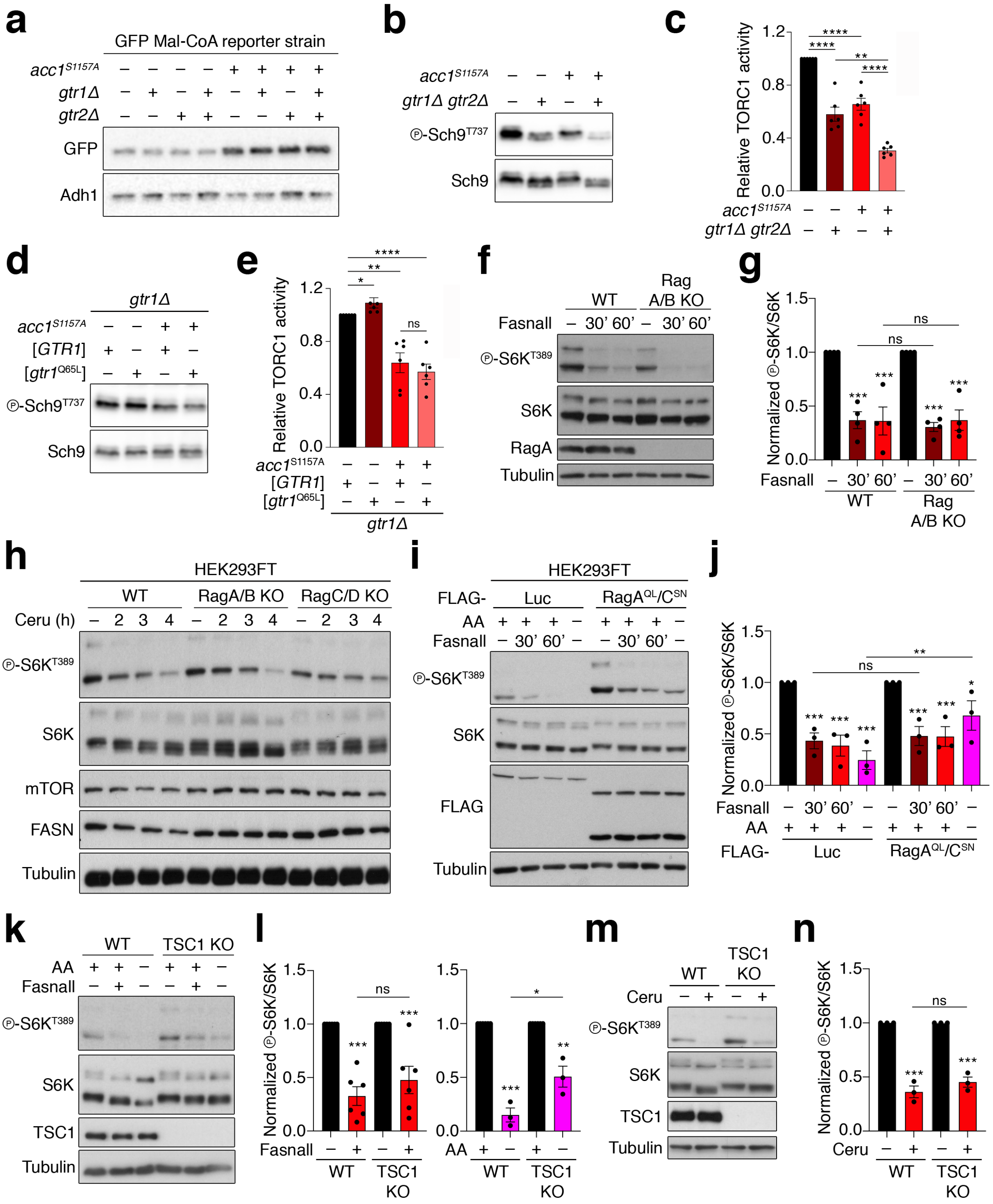


**Extended Data Figure 7. Perturbations in FASN and Acc1 affect mTORC1 activity independently from the Rag and TSC complexes.**

**(a)** Individual or combined loss of Gtr1 and Gtr2, alone or in combination with *acc1*$^{S1157A}$, does not influence Mal-CoA levels. Immunoblots with lysates from the indicated yeast strains, also expressing the fapR/fapOp-yeGFP reporter system, analysed as in Fig. 1b. Mal-CoA levels indicated by GFP expression.

**(b-c)** Expression of the *acc1*$^{S1157A}$ allele inhibits TORC1 in exponentially growing cells in a Gtr-independent manner. Immunoblots with lysates from the indicated yeast strains assayed for TORC1 activity as in Fig. 1b (b). Quantification of TORC1 activity (p-Sch9$^{T737}$/Sch9) in (c), n = 6.

**(d-e)** Expression of the GTP-locked Gtr1$^{Q65L}$ allele does not suppress the TORC1 inhibition mediated by the Acc1$^{S1157A}$ allele. The indicated yeast strains were grown exponentially and assayed for TORC1 activity as in Fig. 1b. Plasmid-encoded genes shown in brackets (d). Quantification of TORC1 activity (p-Sch9$^{T737}$/Sch9) in (e), n = 6.

**(f-g)** FASN inhibition downregulates mTORC1 activity independently from the Rags. WT or RagA/B KO HEK293FT cells were treated with 25 μM Fasnall for the indicated times and mTORC1 activity was assayed by immunoblotting (f). Quantification of mTORC1 activity (p-S6K$^{S389}$/S6K), normalized to each DMSO-treated control, in (g), n = 4.

**(h)** WT, RagA/B KO, or RagC/D KO HEK293FT cells were treated with 50 μM cerulenin (Ceru) for the indicated times and mTORC1 activity was assayed by immunoblotting and S6K phosphorylation (f). n = 5.

**(i-j)** Constitutively active Rags that blunt the AA starvation response do not prevent mTORC1 downregulation by FASN inhibition. HEK293FT cells were transiently transfected with vectors expressing FLAG-tagged RagA$^{QL}$ and RagC$^{SN}$ mutants, or Luciferase (Luc) as control, and treated with 25 μM Fasnall for the indicated times or with AA starvation media for 1 h. mTORC1 activity assayed by immunoblotting (i). Quantification of mTORC1 activity (p-S6K$^{S389}$/S6K), normalized to each DMSO-treated control, in (j), n = 3.

**(k-l)** FASN inhibition downregulates mTORC1 activity independently from the TSC. WT or TSC1 KO HEK293FT cells were treated with 25 μM Fasnall (30 min) or with AA starvation media (1 h). mTORC1 activity was assayed by immunoblotting (k). Quantification of mTORC1 activity (p-S6K$^{S389}$/S6K), normalized to each DMSO-treated control, in (l), n = 6 for Fasnall (left), n = 3 for AA starvation (right).

**(m-n)** As in (k-l), but using 50 μM Cerulenin (Ceru) (4 h) to inhibit FASN (m). Quantification of mTORC1 activity (p-S6K$^{S389}$/S6K), normalized to each DMSO-treated control, in (n), n = 3.

Data in all graphs shown as mean ± SEM. * = $p < 0.05$, ** = $p < 0.005$, *** = $p < 0.0005$, **** = $p < 0.00005$.

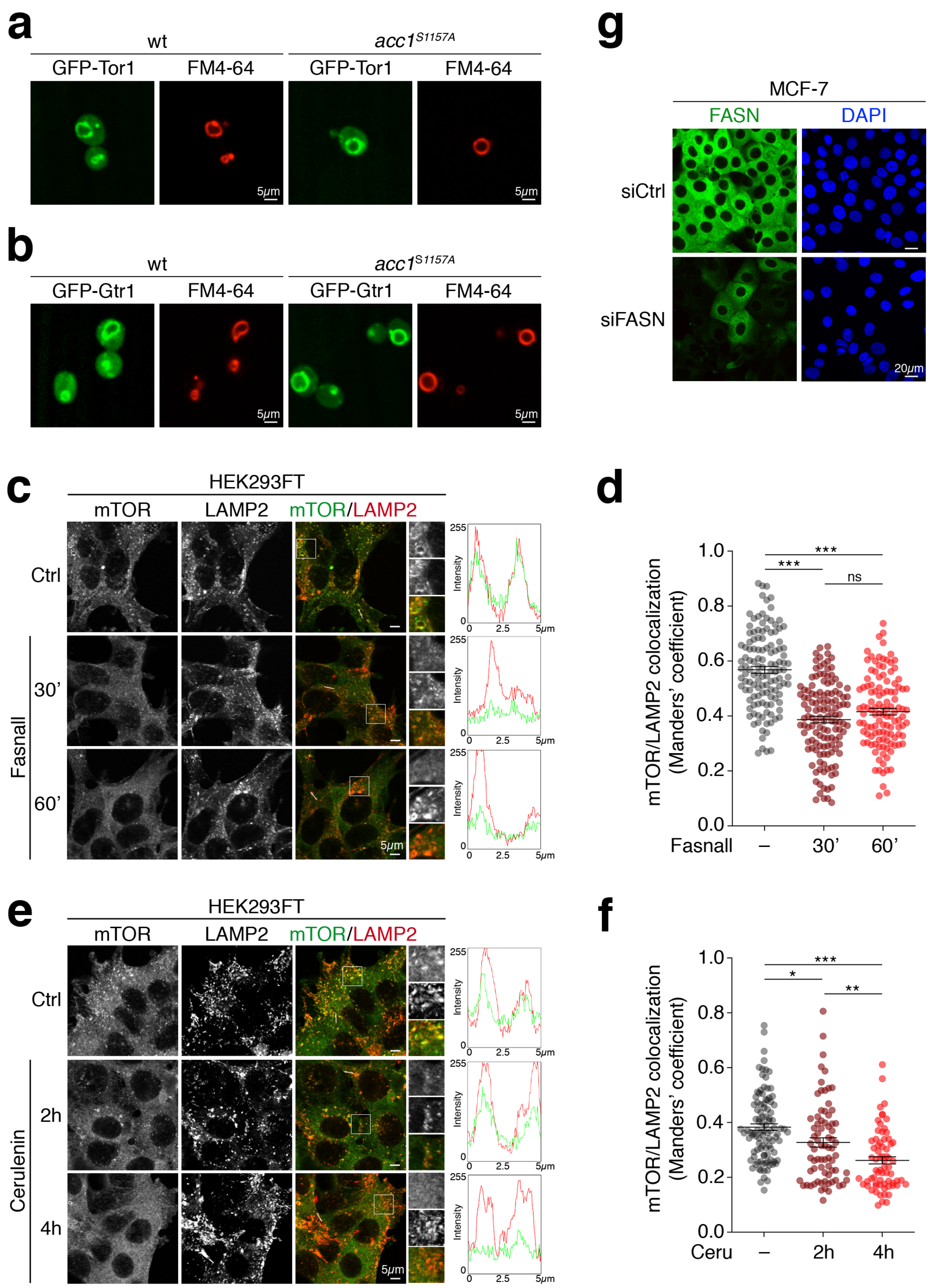


**Extended Data Figure 8. mTOR/Tor1 and Gtr1 localization upon FASN inhibition or yeast Acc1 hyperactivation.**

**(a-b)** Constitutive activation of Acc1 does not alter vacuolar morphology or the subcellular localization of GFP-Tor1 (a) or GFP-Gtr1 (b). Wild-type (wt) or *acc1*$^{S1157A}$ cells expressing GFP-Tor1 or GFP-Gtr1 from plasmids were grown exponentially and localization of GFP fusion proteins was assayed by fluorescence microscopy. Vacuoles stained with the vacuolar membrane fluorescent dye FM4-64. Scale bars = 5 µm.

**(c-d)** Colocalization analysis of mTOR with LAMP2 (lysosomal marker) in HEK293FT cells treated with 25µM Fasnall for the indicated times using confocal microscopy. Magnified insets and intensity plots for selected regions shown to the right. Scale bars = 5 µm (c). Quantification of colocalization from individual cells in (d). Pooled data from n = 4 independent experiments.

**(e-f)** As in (c-d), but inhibiting FASN with 50 µM cerulenin for the indicated times. Scale bars = 5 µm (e). Quantification from individual cells in (f). Pooled data from n = 3 independent experiments.

**(g)** Specific staining of endogenous FASN by immunofluorescence. Control or *FASN* knockdown MCF-7 cells were stained with an anti-FASN antibody and signal from endogenous FASN was detected by confocal microscopy. Nuclei stained with DAPI. Scale bars = 20 µm.

Data in dot plots shown as mean ± SEM. * = $p < 0.05$, ** = $p < 0.005$, *** = $p < 0.0005$.

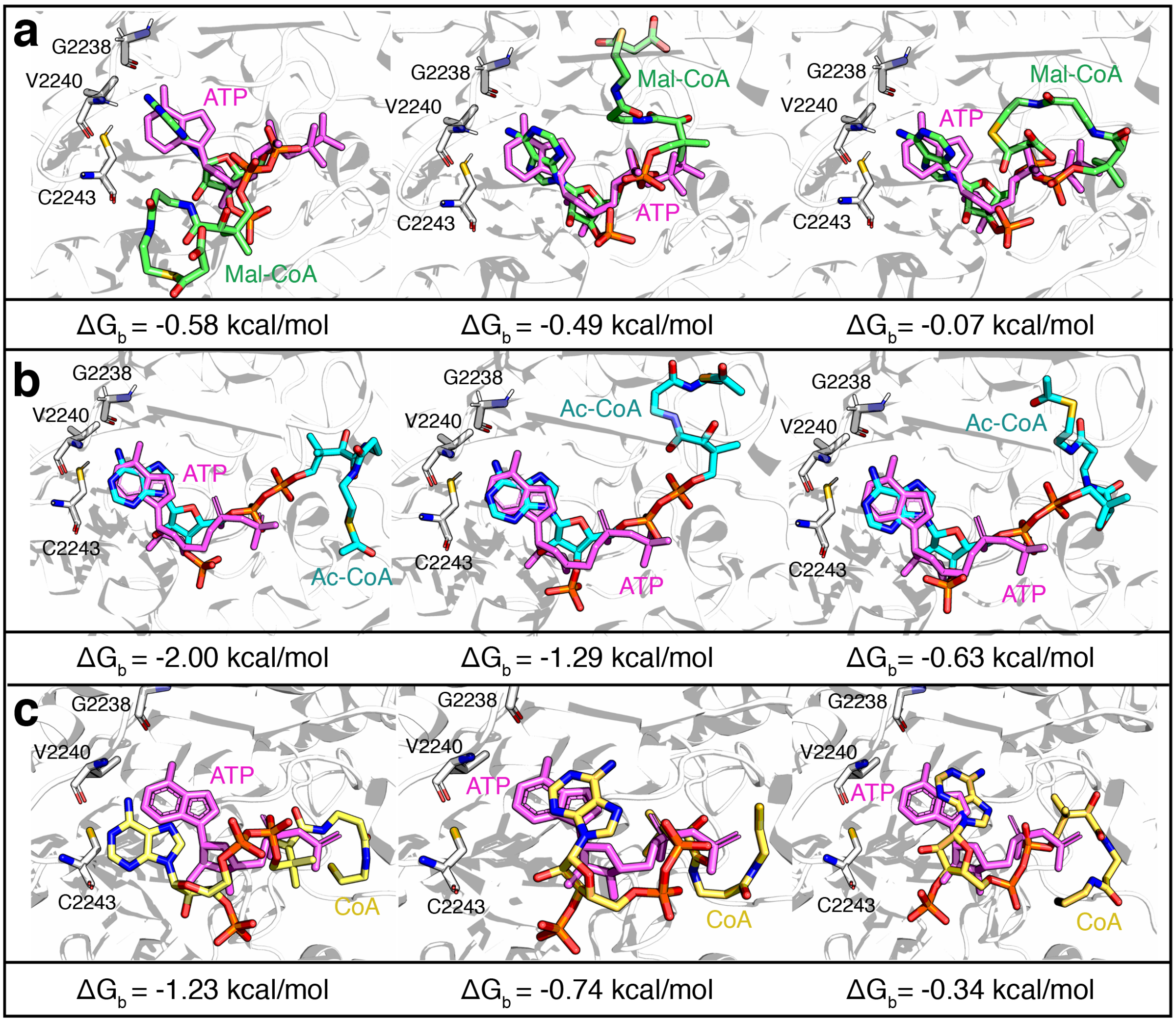


**Extended Data Figure 9. Site-specific docking simulations for Mal-CoA, Ac-CoA, and CoA in the catalytic pocket of mTOR.**

**(a)** The best three docking poses of malonyl-CoA (Mal-CoA; shown in green) aligned to ATP (violet) in the mTOR catalytic pocket. The binding energy value ($\Delta G_b$) for each conformation is shown below the docking models. These conformations were selected as the starting point to perform all atom MD simulations.

**(b)** As in (a), but for acetyl-CoA (Ac-CoA), shown in cyan.

**(c)** As in (a), but for Coenzyme A (CoA), shown in yellow.

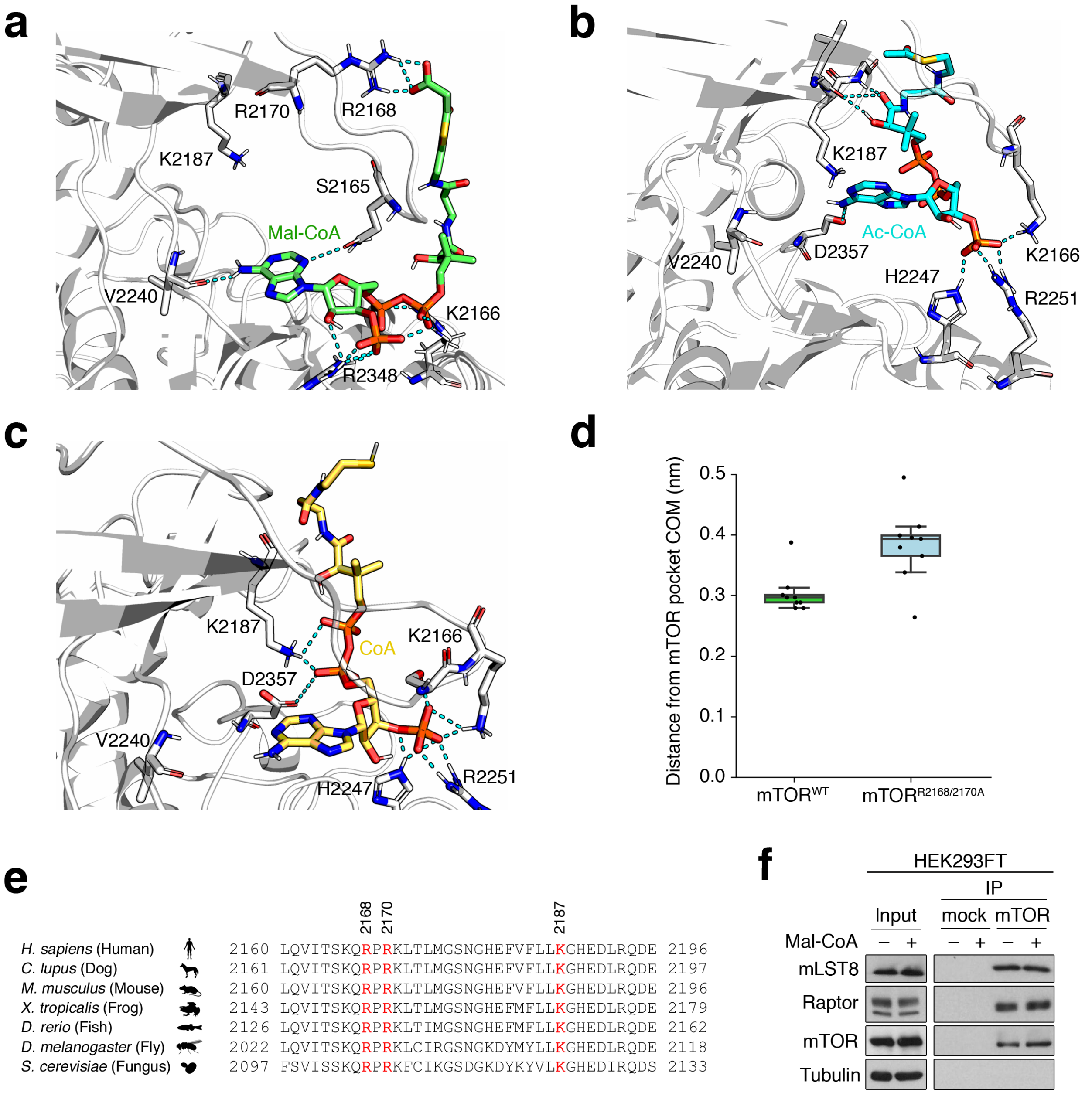


**Extended Data Figure 10. Molecular dynamics simulation and docking experiments suggest that charged interactions stabilize Mal-CoA in the mTOR pocket.**

**(a)** The malonyl group of Mal-CoA forms salt bridges with charged residues at the edge of the mTOR catalytic pocket (lateral view). Representative Mal-CoA (green) conformation sampled by MD simulations. The hydrogen bonds established between Mal-CoA (final conformation in the simulation) and the amino acid residues of the mTOR pocket are shown as cyan dotted lines. Note that only a snapshot is shown, with multiple residues participating in the formation of dynamic interactions with the malonyl group, with R2168 being the most frequent.

**(b-c)** Representative acetyl-CoA (Ac-CoA) placement in the mTOR binding pocket (b). Representative CoA conformation sampled by MD simulations in (c). Hydrogen bonds between the compounds and the amino acid residues of the mTOR catalytic pocket indicated by cyan dotted lines.

**(d)** *In silico* mutagenesis of key mTOR residues weakens Mal-CoA interactions with mTOR. Distances of Mal-CoA from the mTOR binding pocket during the MD simulations as in Fig. 3c, comparing $mTOR^{WT}$ and $mTOR^{R2168A/R2170A}$ molecules. Three replicas were run, with individual dots representing averages over 100 ns of MD simulation (n = 9). Data in box plots: central line, median; box, interquartile range (IQR) [$25^{th}$ (Q1) - $75^{th}$ (Q3) percentile]; whiskers, Q3+1.5*IQR and Q1-1.5*IQR.

**(e)** Amino acid sequence alignment of the 2160-2196 aa region of human mTOR with the respective orthologous sequences from other organisms. Key conserved residues that participate in interactions with Mal-CoA shown in red.

**(f)** Mal-CoA does not affect mTORC1 complex stability. HEK293FT cells were lysed and endogenous mTOR was immunoprecipitated in the presence or absence of 1 mM Mal-CoA (added directly in the lysates 5 min prior to addition of the antibody). Co-immunoprecipitation of the mTORC1 subunits Raptor and mLST8 assayed by immunoblotting.
